# Neurometabolic predictors of mental effort in the frontal cortex

**DOI:** 10.1101/2024.01.23.576854

**Authors:** Arthur Barakat, Jules Brochard, Mathias Pessiglione, Jean-Philippe Godin, Bernard Cuenoud, Lijing Xin, Nicolas Clairis, Carmen Sandi

**Author notes:** Corresponding & lead author. Co-senior author. Psychiatry Department, Faculty of Medicine, University of Geneva, Geneva, Switzerland (Nicolas Clairis).

## Abstract

Motivation drives individuals to overcome costs to achieve desired outcomes, such as rewards or avoidance of punishment, with significant variability across individuals. The dorsomedial prefrontal cortex/dorsal anterior cingulate cortex (dmPFC/dACC) and anterior insula are key brain regions implicated in effort-based decision-making. Here, we utilized proton magnetic resonance spectroscopy (^1^H-MRS) at 7 Tesla on 69 healthy participants in these brain regions to uncover the neurometabolic factors that influence these differences. We designed and applied an effort-based decision-making task requiring mental and physical effort to probe motivated behavior, complemented by computational modeling to extract key behavioral parameters. Gradient boosting machine learning was applied to explore the predictive role of specific metabolites in motivated behavior. Our results reveal that a model established on dmPFC/dACC metabolites explains decisions to exert high mental effort and sensitivity to mental effort. In particular, glutamate, aspartate, and lactate in dmPFC/dACC, three metabolites linked through the tricarboxylic acid cycle and glycolysis, were identified as key discriminative metabolites in the dmPFC/dACC, predictive of mental effort choices, underpinning energy supply and cognitive processes. Anterior insula metabolites did not significantly relate to effort-related decisions. Notably, glutamine and lactate levels between the periphery (plasma) and the dmPFC/dACC were correlated, suggesting a metabolic link between peripheral and central biomarkers of effort. Our findings provide novel insights into the neurometabolic underpinnings of motivated behavior and propose novel biomarkers for mental effort-based decision-making. Importantly, our study highlights the relevance of multivariable approaches in elucidating complex cognitive functions.

## Introduction

Motivation drives individuals to overcome the inherent costs of actions to achieve desired outcomes, such as obtaining rewards or avoiding punishments ^1,2^. This process, fundamental to human behavior, involves making decisions based on cost-benefit trade-offs between the rewards involved in an action and the effort required to reach them. However, there is striking variability in how individuals approach effortful decisions. These inter-individual differences can significantly impact well-being, longevity, and success in life ^3–5^. Moreover, a pronounced aversion to effort exertion is a key symptom in various brain pathologies ^2,6^. While such diversity in motivational drive reflects a significant aspect of human cognition, our understanding of the neurobiological underpinnings of these inter-individual differences remains limited.

Neurobiologically, motivation and effort-based decision-making are linked to the functioning of the dorsomedial prefrontal cortex/dorsal anterior cingulate cortex (dmPFC/dACC) ^2,7–10^ which plays a pivotal role in driving mental and physical effort ^11–13^. Lesions in this area can lead to increased effort aversion ^14^ and even akinetic mutism ^15^, a state characterized by a lack of initiative to act. However, our understanding of the specific neurobiological components within the dmPFC/dACC that influence effort-based decision-making is still limited.

Recent advances emphasize the importance of brain bioenergetic and metabolic processes for brain function, behavior, and cognition ^16–18^. It is therefore essential to understand whether the dmPFC/dACC baseline metabolism also influences effort-based decision-making. Initial studies interested in the link between brain metabolism and cognitive effort predominantly focused on glucose as a resource-limiting energy source for demanding cognitive processes ^19,20^. However, brain glucose does not seem to be the resource-limiting energy resource for cognitive processes that was originally proposed ^21–24^. Additionally, neurons utilize other energy substrates to support both neural structure and function ^25–29^ and, consequently, meet cognitive demands ^18^. Several animal and human studies suggest that neurometabolic components in different brain regions are involved in decision-making and motivation ^30–36^. However, only a few studies focused on the relationship between dmPFC/dACC metabolism and cognitive effort yet ^37–39^ and most of these studies investigated cognitive performance rather than the motivation to engage with cognitively effortful tasks. It is therefore important to study the link between dmPFC/dACC metabolism and motivated behavior towards cognitively effortful tasks.

Brain metabolism can be studied with magnetic resonance spectroscopy (MRS) which enables the non-invasive quantification of brain metabolites thereby providing insights into the neurochemical state of specific brain regions. Proton (^1^H)-MRS enables the quantification of several brain metabolites^40^, including glutamate, lactate, aspartate, N-acetylaspartate (NAA) or creatine. These metabolites play crucial roles in neuronal health, energy metabolism, and cellular signaling, all key for neural function and behavior production ^40–42^. Understanding whether the concentrations of these metabolites in the dmPFC/dACC are related to effort-based motivated processes can provide invaluable insights into the neurometabolic foundations of these cognitive functions. However, virtually all the studies linking metabolite concentrations, acquired through ^1^H-MRS, to behavior or cognition have focused on individual metabolites using univariate analyses, overlooking how the combined influence of multiple metabolites contributes to motivated behavior, despite the fact that most metabolites share and interact within overlapping metabolic pathways ^42^. Addressing this gap by employing multivariable statistical methods can provide a more comprehensive view of the dmPFC/dACC neurochemical bases of motivated behavior.

Additionally, the anterior insula (AI) is also often co-activated with the dmPFC/dACC with which it forms the salience network ^43^. Accordingly, it is also involved in mental and physical effort ^2,11,44^. It is therefore of utmost importance to understand whether the metabolism of these two effort nodes equivalently predicts effort-based motivation.

In this study, we leveraged the critical role of the dmPFC/dACC in assessing cost-benefit trade-offs to explain individual differences in decision-making for effort-based motivated behavior. We utilized ultra-high field 7 Tesla (7T) ^1^H-MRS to measure metabolite concentrations in the dmPFC/dACC and in the anterior insula (AI). Additionally, we conducted metabolomic analyses of plasma samples. To gauge participants’ willingness to exert physical and mental efforts, we created an effort-based decision-making task where subjects chose between a fixed low effort/low incentive option and a varying high effort/high incentive option. The task involved trials in which participants could obtain rewards or avoid punishment, alternating between mental and physical effort blocks. We applied computational modeling to extract key parameters influencing effortful decisions in our task, such as mental and physical effort perception ^2,45^. We subsequently employed machine learning models to explore whether specific dmPFC/dACC and AI metabolite combinations could predict effortful choices and parameters of motivated behavior. Our study offers new insights into how the dmPFC/dACC metabolic processes relate to motivated behavior components, potentially identifying novel biomarkers and therapeutic targets for motivation-related cognitive functions, especially in the context of mental effort and perception.

## Methods

### Participants

To ensure sufficient statistical power and heterogeneity of the input features for machine learning model training and evaluation, a total of 75 healthy volunteers (40 females from which 11 reported using a hormonal contraceptive method) participated in the study, approved by the Cantonal Ethics Committee of Vaud (CER-VD), Switzerland. All methods were performed in accordance with the relevant guidelines and regulations. Data analyzed here is part of a larger study from which a report focusing on physical effort has been previously published [36]. Exclusion criteria included being outside the 25 - 40 years age range, regular drug or medication use, history of neurological disorders, and MRI contraindications. Informed consent was obtained from all participants before being invited to the study. Participants also completed several online questionnaires before their visit in the laboratory, including the Montgomery Asberg Depression Rating Scale – self-rated (MADRS-S). To ensure sufficient inter-individual variability in motivation, we stratified recruitment based on the MADRS-S, with a cutoff of 4 differentiating non-depressive (N = 37) from mild/high depressive (N = 38) individuals ^46^. After exclusion for incomplete or incongruent behavior, the sample included 71 subjects (35 females; mean age = 30.2 ± 3.4 years). See Supplementary Methods for further details.

### Experimental procedure

Participants were requested to abstain from eating in the hour preceding their visit in the laboratory at 2pm. Blood samples were collected by healthcare professionals in dedicated facilities (the EPFL “Point Santé”, i.e. EPFL infirmary, or the Arcades medical center, both located in Lausanne). Immediately after, participants were brought to the MRI facility where baseline metabolite concentration assessments were performed in the dmPFC/dACC and AI using proton magnetic resonance spectroscopy (^1^H-MRS) (**Fig. 1a-c**). Participants then received instructions about the preliminary training outside of the scanner and the behavioral task to be performed during fMRI acquistion.

**Fig. 1:**
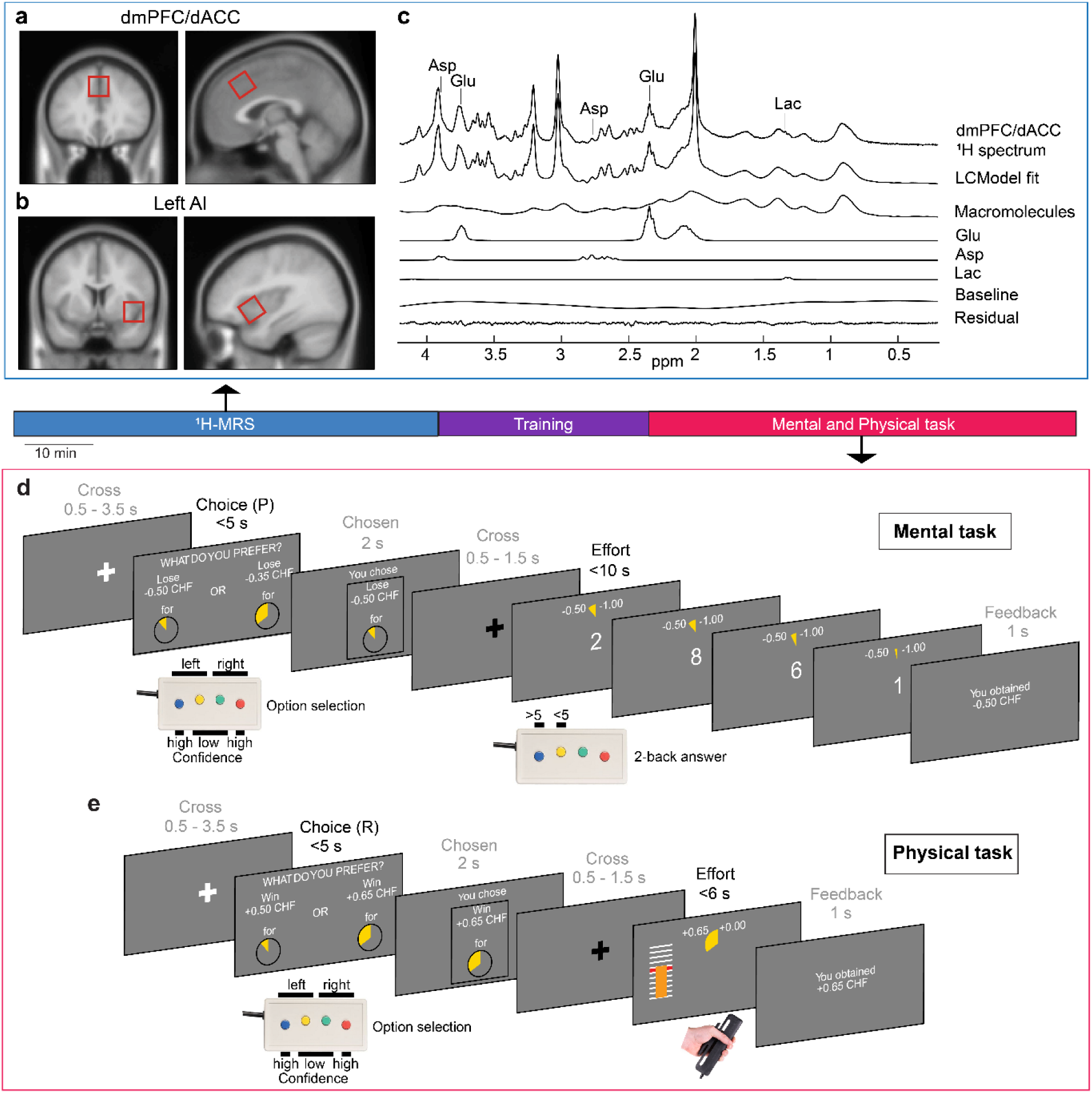
Overview of experimental design. (**a-b**) Localization of dmPFC/dACC and AI voxels (20 x 20 x 20mm³) targeted for ^1^H-MRS acquisitions. The voxels are displayed in coronal and sagittal sections, overlaid on the MNI152 T1 average anatomical scan for reference. (**c**) Example of a ^1^H-MR spectrum obtained from the dmPFC/dACC. The spectrum showcases the LGCModel spectral fit, macromolecular signals, baseline, residual fit, and individual metabolite fits for glutamate (Glu), aspartate (Asp), and lactate (Lac). (**d-e**) Effort-based decision-making tasks. Participants choose between options varying in effort and monetary incentive (reward or punishment) levels, with the potential for higher effort leading to greater rewards or avoiding larger punishments. Each trial started with a fixation cross, followed by a 5-seconds window where participants had to pick up the option that looked the more interesting for them based on their subjective preferences. There was always one option associated to a low effort and a low monetary incentive, while the other was associated to a larger monetary incentive but also to a higher effort. Physical and mental efforts were segregated into distinct blocks (2 for each type of effort), while rewards and punishment incentives were intermixed across trials within each block. After the choice was made, the chosen option was briefly displayed, followed by another fixation cross with a jitter to separate decision-making from the effort phase. (**d**) In the mental effort task, participants aimed to achieve a target number of correct responses within 10 seconds, determined by their prior calibration and the selected effort level. (**e**) In the physical effort task, participants had to maintain force above 55% of their maximal force for a duration depending on the chosen effort level (between 0.5 and 4.5s). Each trial ended with performance feedback indicating how much money was earned (or lost respectively) in the current trial.

### Behavioral task

The behavioral task was designed and run using Psychtoolbox 3 ^47^, and implemented in Matlab (The Mathworks Inc., USA). The main task aimed at characterizing inter-individual differences in the motivation to engage with physical or mental effort across individuals via a binary choice task (**Fig. 1d**). Motivation depends on a cost/benefit tradeoff between the expected benefits of the goal to be reached (reward to obtain/loss to avoid) and the expected costs (e.g., physical/mental effort) to reach that goal ^2,48^. We therefore designed a task to capture these different components of motivated behavior. The task comprised both positive (monetary gain) and negative (monetary loss) valence trials to account for these two types of motivational incentives. Physical and mental effort were presented in different blocks both during the training and during the main task. The main task consisted in selecting between one low incentive (low monetary gain (+0.5 CHF), or high monetary loss (−0.5 CHF) in the loss domain respectively)/low effort (effort level E0) option and one high incentive (high monetary gain, or low monetary loss in the loss domain respectively)/high effort option on each trial. Gain incentives (R1/R2/R3), loss incentives (P1/P2/P3) and effort levels (E1/E2/E3) associated with the high incentive/high effort option varied independently between 3 levels each based on the calibration (see **Supplementary Methods** for more details). Incentive levels were calibrated by identifying an indifference point for each effort type during the training period. The participants made 216 choices between one low incentive/low effort and one high incentive/high effort option (left/right position on the screen counterbalanced across trials). While selecting an option, participants were also requested to provide a confidence rating on their choice (defined as the subjective probability of having selected the best option) as each option was associated with either a low or a high confidence button (**Fig. 1d**). This confidence rating aimed at improving the quality of the computational modeling applied to the choices. After their choice, participants had to perform the effort chosen to obtain the corresponding monetary gain (or avoid doubling the corresponding monetary loss). Physical effort implied to exert at least 55 % of an individual’s maximum voluntary contraction force (MVC), which had been previously calibrated during the training period, during a varying amount of time (0.5-4.5s) depending on the chosen effort. Mental effort implied to provide a certain number of correct answers in a 2-back task according to the maximum number of correct responses (MNCR) which had been previously calibrated during the training period, and to the effort chosen. Physical and mental effort were presented in blocks of 54 trials alternating with each other in a counterbalanced order across individuals. The complete set of explanations is reported in the **Supplementary Methods**.

### Data & Code availability

The original code developed for the task has been deposited at https://github.com/NicolasClairis/physical_and_mental_E_task.git and is publicly available as of the date of publication. The data and code to reproduce the analyses of the paper is available from the lead contact upon request.

### MRS acquisition and preprocessing

Proton magnetic resonance (^1^H-MR) dmPFC/dACC and AI spectra (**Fig. 1c**) were collected using a 7 Tesla/68 cm MR scanner (Magnetom, Siemens Medical Solutions, Erlangen, Germany) with a single-channel quadrature transmitter and a 32-channel receive coil (Nova Medical Inc., MA, USA). See

**Supplementary Methods** for details on the ^1^H-MRS acquisition and analysis and **Figure S1** for the average location of the ^1^H-MRS voxels across participants.

### Computational modeling of motivated behavior

Participants’ choices were fitted with a softmax model using Matlab’s VBA toolbox (https://mbb-team.github.io/VBA-toolbox/) which implements Variational Bayesian analysis under the Laplace approximation ^49^. The details of the model have been described in our previous publication ^36^, but the details can also be found in the **Supplementary Methods**. In brief, the model fits the probability of choosing the high effort option with a sigmoid transformation aiming at extracting individual parameters reflecting the weight that different task parameters had on decision-making. These parameters included sensitivity to reward (kR), punishment (kP), physical effort (kEp), mental effort (kEm), physical fatigue (kFp), mental learning (kLm) and a bias term (bias) capturing the general propensity to engage with the task. Subsequently, all parameters except the bias term were boxcox transformed to reduce skewness and increase normality ^50^.

### Feature selection and extreme gradient boosting trees (XGBoost)

Our main goal was to test whether we could predict the proportion of high physical effort (HPE) (model 1) and/or the proportion of high mental effort (HME) (model 2) choices based on the brain metabolites using two machine learning models. Subsequently, we also developed a machine learning model (model 3) based on the same brain metabolites to predict the sensitivity to mental effort (kEm) derived from our computational model. Finally, we tried to reduce the number of metabolites used for the prediction of HME to identify a subset of metabolites that are sufficient to significantly predict the proportion of HME choices (model 4). Due to the fact that some participants did not complete all the trials of the mental effort task, we had to remove 2 additional subjects from the analysis of HME (models 2 and 4) leading to a sample size of 69 subjects (34 females; mean age = 30.5 ± 3.9 years; mean BMI = 22.8 ± 3.1) for these models. However, since this did not impair the estimation of the behavioral parameters, model 3 was still performed on the 71 study subjects. Feature selection plays a crucial role in developing predictive models, as it simplifies the model by removing redundant features and mitigates overfitting, particularly for small sample sizes with limited generalization ability. Our dataset was first randomly divided in a train/test split, with 80% of the dataset kept for training and 20% for testing. However, to increase the robustness of our results for kEm predicions we increase the train/test split, with 75% of the dataset kept for training and 25% for testing. To avoid bias in test predictions, feature selection was conducted on the training dataset for both models. Features with a Pearson correlation coefficient smaller than 0.1 with the target variable were removed. As a second round of feature selection, Sklearn.*AdaBoostRegressor* package was used to train models using approach CVLOO to predict target variables. The selected features were those that yielded the smallest validation error from the CVLOO approach. The resulting metabolites were used to train both machine learning algorithms.

We employed *XGBoost* to predict the number of high physical effort (HPE) and high mental effort (HME) choices selected by the participants (models 1, 2 and 4) and the mental effort perception kEm (model 3), extracted with the behavioral computational model, using a selected set of metabolite concentrations. We used the *XGBRegressor* function from the Python *XGBoost* package to fit both models. XGBoost includes several adjustable hyperparameters. We optimized the step size shrinkage (eta), maximum depth of the tree (max_depth), minimum sum of instance weight (min_child_weight), and regularization parameters (gamma, lambda, alpha) through Bayesian optimization with the *hyperopt* Python package and CVLOO ^51^. We assessed our model’s prediction performance by correlating the predicted and true values of the target variables in a holdout sample that was not utilized in the model learning process. The train and test datasets were loaded into data frames using the *pandas* Python package and evaluated using model assessment metrics computed with the *numpy* Python package. Additionally, we computed the Shapley Additive exPlanations (SHAP) values using the *shap* Python package ^52^.

## Results

Our study was designed to explore how inter-individual variation in 7T proton magnetic resonance spectroscopy (^1^H-MRS) measured metabolites in the dmPFC/dACC (**Fig. 1a,c**) can predict differences in physical and mental motivated behavior across individuals. Additionally, we performed ^1^H-MRS scans in the AI to assess the specificity of our predictive findings for the dmPFC/dACC or the network constituted by the dmPFC/dACC and the AI (**Fig. 1b**), as AI activity often co-varies with the dmPFC/dACC within the salience network ^43^ and is also related to effort ^2,11,44^. To minimize any external influences (digestion, circadian rhythms, etc.) on the metabolic state of our participants, we always started the experiment at 2 PM and participants were asked to avoid eating at least 1h before. Participants also had to avoid from performing any intense physical activity on the day of the experiment and on the preceding day.

### Behavioral task

To assess motivated behavior, participants performed a behavioral task comprising 216 choice trials divided into 4 blocks of 54 trials. In each trial, participants had to choose between a low monetary incentive associated with a fixed low effort option and a varying high monetary incentive/high effort option (**Fig. 1d-e**). They subsequently had to execute the chosen effort on each trial to ensure that their decisions accurately reflected the presented effort levels. The task was divided between 2 mental effort blocks (2-back task) and 2 physical effort (squeezing a handgrip) blocks alternating with each other in a counterbalanced order across individuals. To ensure task feasibility and prevent participants from perceiving decisions as risky which could bias the effort perception ^53^, the task was calibrated to each individual’s performance during the training period (**Fig. 1**). Monetary incentives included both gain and loss trials to account for a potential ‘loss aversion’ bias ^54^, which implies a higher motivation for minimizing monetary losses than for maximizing monetary gains. Notably, participants successfully completed the chosen effort in 95 ± 3.7% (mean ± SD) of the trials involving mental effort, and in 98 ± 3.4% of those requiring physical effort confirming that participants were capable of performing the effort chosen (i.e., the influence of risk on decision-making was minimized). Increased effort difficulty, associated with the high-effort option, resulted in fewer selections of high-effort/high-reward incentives in both physical and mental tasks **(Fig. S2a,d**), indicative of effort aversion. Effort aversion can be explained by the fact that, in our task, selecting higher difficulty levels led to tangible consequences: in the physical task, choosing the more effortful option correlated with higher handgrip force production (**Fig. S2b-c**). In the mental task, it resulted in a higher number of errors that seemed compensated by an elevated cognitive efficiency, defined as the ratio between the number of correct answers divided by time (**Fig. S2e-f**). In other words, these outcomes corroborate the task’s effectiveness in quantifying motivational dynamics for both types of effort.

Importantly, the proportion of high mental effort choices and high physical effort choices were very weakly correlated to one another (r = 0.22, p = 0.06), confirming that they could be observed as the results of at least partially independent processes.

### dmPFC/dACC metabolites explain motivated behavior

To determine if dmPFC/dACC metabolite levels can predict the proportion of high mental effort (HME) and high physical effort (HPE) effort choices made by participants, we employed a gradient tree boosting regression model with metabolite concentrations as regressors. Our feature selection process, applied to 18 features including metabolites and standard ratios (detailed in **Supplementary Materials**), identified 8 features relevant for HME and 4 for HPE prediction (**Fig. 2**). To enhance data robustness and minimize overfitting, we adopted a train/validation/test approach with cross-validation leave-one-out (CVLOO) design, randomly splitting data into training/validation and testing sets with an 80% ratio, resulting in training/validation (N = 55) and testing (N = 14) datasets. We trained extreme gradient boosting trees (XGBoost) to fit linear response functions to HME and HPE, utilizing Bayesian optimization for hyperparameter tuning. The prediction error of the model was quantified using the root mean square error (RMSE).

**Fig. 2:**
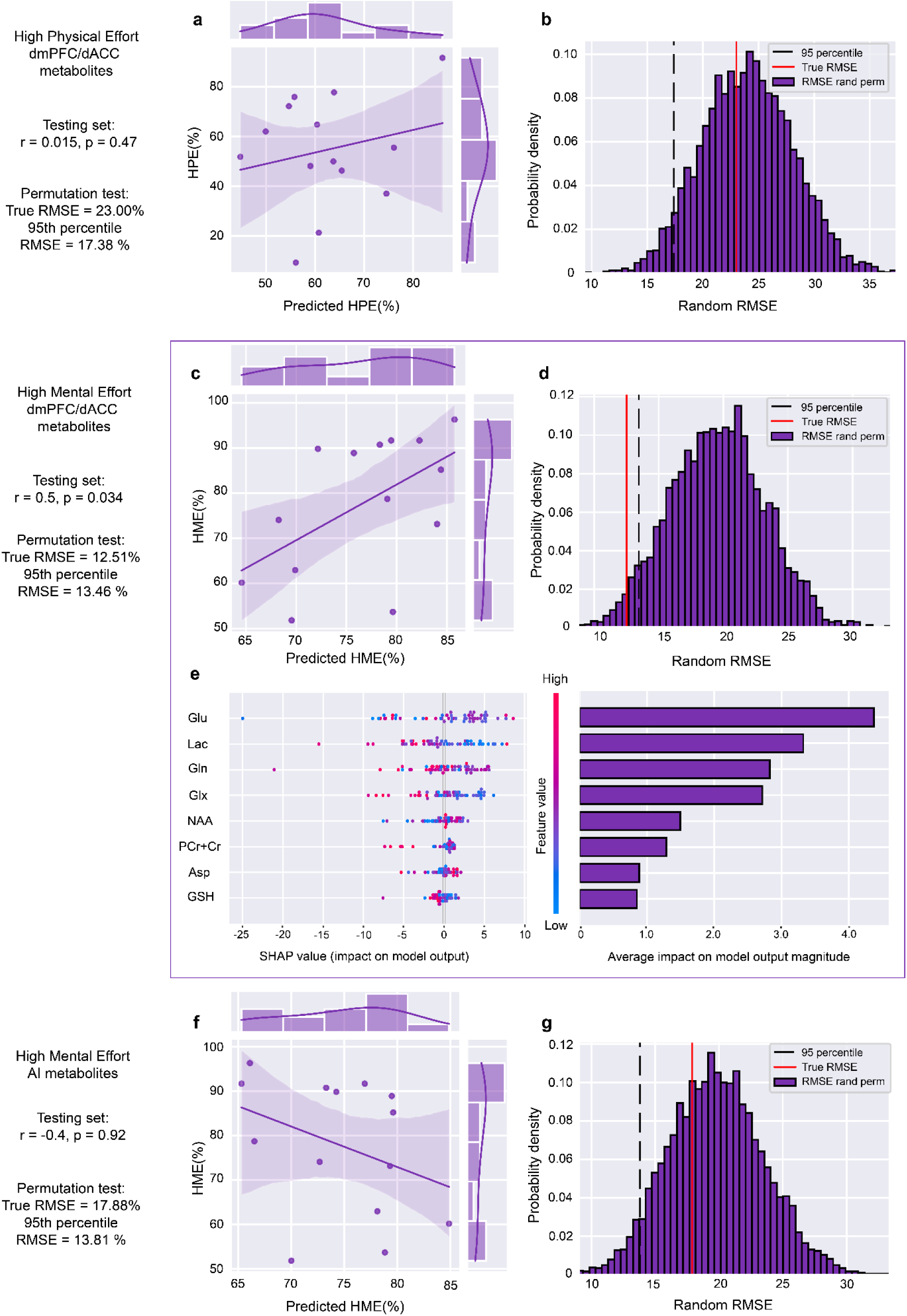
Analysis of models’ explanations for physical and mental effort choices based on metabolite concentrations in dmPFC/dACC and AI. (**a**) Analysis of high physical effort (HPE) explanations using dmPFC/dACC metabolite concentrations with an XGBoost model (model 1) indicates a moderate correlation not reaching significance (r = 0.015, p = 0.47) for the test set. (**b**) Furthermore, a permutation test indicated lack of predictive robustness (95^th^ percentile threshold = 17.38% vs. model RMSE = 23.00). (**c**), For high mental effort (HME) choices (model 2), our model showed a strong correlation (r = 0.5, p = 0.034), underscoring the model’s efficacy in anticipating decision patterns based on dmPFC/dACC metabolite profiles. (**d**) Permutation test for HME predictions confirms model accuracy (95^th^ percentile threshold = 13.46% vs. model RMSE = 12.51%). (**e**) Feature importance analysis via SHAP values for HME prediction identifies nine key metabolites: glutamate (Glu), lactate (Lac), glutamine (Gln), glutamate + glutamine (Glx), N-acetylaspartate (NAA), N-acetylaspartylglutamate (NAAG), aspartate (Asp), glutathione (GSH), and phosphocreatine+creatine (PCr + Cr). In brief, the color code corresponds to each metabolite’s concentration and the SHAP value indicates a more positive or a more negative impact on the outcome variable. (**f**) A model trained on AI metabolites’ predictive power for HME reveals a negative correlation between predicted and actual HME that does not reach statistical significance (r = -0.4, p = 0.92). (**g**), A permutation test further indicates a significant deviation in model performance for AI-based predictions (95^th^ percentile threshold = 13.81% vs. model RMSE = 17.88%), highlighting a failure to predict HME choices based on AI metabolites.

For HPE, model 1 showed a medium fit in training and validation (train RMSE = 0.16%, validation RMSE = 16.06%), and a non-significant result on the test set (test RMSE = 23.00%, r = 0.015, p = 0.47), explaining only 5% of the variance (**Fig. 2a**). A permutation test with 5000 permutations further indicated that these results were not above chance level (95^th^ percentile = 17.38 > model RMSE = 23.00) (**Fig. 2b**), indicating that HPE could not be significantly predicted with the combinations of baseline dmPFC/dACC metabolites available in our dataset. Similarly, training a model on metabolites from the AI, resulted in poor and non-significant performance (test RMSE = 24.46%, r = 0.24, p = 0.4), leading us to discard HPE prediction from further analysis.

In contrast, for HME, the model 2 demonstrated a good fit (train RMSE = 0.05%, validation RMSE = 13.05%) and a consistent result in the unbiased estimate that is the test set (test RMSE = 12.51%, r = 0.49, p = 0.034), explaining 25% of the variance (**Fig. 2c**). The permutation test with 5000 permutations confirmed that our model’s predictions significantly exceeded chance level (95^th^ percentile = 13.48 > model RMSE = 12.51) (**Fig. 2d**). These results support our hypothesis that dmPFC/dACC metabolite concentrations can predict inter-individual variation in motivated behavior, specifically demonstrating their predictive power in the context of mental effort tasks.

To understand the impact of individual metabolites in the XGBoost model for predicting HME, we analyzed the Shapley Additive exPlanations (SHAP) values. The identified features included in descending order of their importance (determined by the magnitude of each feature’s SHAP value) are glutamate (Glu), lactate (Lac), glutamine (Gln), Glutamine + Glutamate (Glx), N-acetylaspartate (NAA), phosphocreatine + creatine (PCr + Cr), aspartate (Asp) and glutathione (GSH) (**Fig. 2e**). Thus, dmPFC/dACC glutamate and lactate levels emerged as the top discriminating features for predicting the motivation to cognitive effort.

Importantly, our SHAP analysis of the data distribution for each predictive feature/metabolite revealed that specific metabolite concentrations do not have a linear relationship with HME (**Fig. 2e**). Indeed, neither glutamate (r = -0.19, p = 0.12) nor lactate (r = -0.13, p = 0.27) displayed a significant linear relationship with HME. Particularly for glutamate, extreme values – either low or high – were negatively associated with HME, indicating a non-linear relationship (**Fig. S3**). Employing Bayesian model comparison to determine the best model linking glutamate with HME, the most accurate fit featured an inverted U-shape relation, based on the mean-centered squared score of glutamate concentrations (r = 0.36, p = 0.003), as supported by both the Akaike Information Criterion (AIC) and Bayesian Information Criterion (BIC) (**Fig. S3a**). However, we could not identify any pattern for lactate’s relationship with HME, supporting that its contribution relies in the multivariable model in a non-linear combination with the other metabolites.

To determine if the metabolites relevant for HME are specific to the dmPFC/dACC region, we similarly assessed the ability of AI metabolite concentrations to predict HME. We applied the same model training process used for the dmPFC/dACC to AI metabolite concentrations. However, this AI-based model did not achieve statistical significance in predicting HME in our test set (r = -0.4, p = 0.92) (**Fig. 2f**), nor in the permutation test (95^th^ percentile = 13.81% > model RMSE = 17.88%) (**Fig. 2g**).

### Computational modeling of motivated behavior

Thus far, our analysis has shown the capability of dmPFC/dACC metabolites to predict HME using XGBoost. However, since motivation, and by extension HME, is a complex construct composed of multiple elements like reward sensitivity or effort aversion ^1,2,45^, the specific behavioral components related to metabolism in this brain region remain to be elucidated. We therefore applied a computational model to help deepening our understanding of behavioral mechanisms and uncovering neurobiological correlates otherwise not observable ^2,55,56^. Specifically, we modelled participants’ choices in order to disentangle the different behavioral parameters that could influence HME in our task. These behavioral parameters are the sensitivities to reward (kR) or punishment (kP), to physical (kEp) or mental (kEm) effort, and the impact of physical fatigue (kFp) or mental learning progress over time (kLm), along with a general bias to select the high effort options (bias).

Following best practices in computational modeling ^57^, we first verified whether our behavioral parameters were recoverable using simulations (**Fig. 3a**) and then confirmed that parameters were not confounded with each other (**Fig. 3b**). Parameter recovery, assessed by comparing simulation parameters with those optimized, was successful in average more than 78% of simulations for all the parameters, except for kLm which was recovered in 44% of simulations, a still rather reasonable result (**Fig. 3a**). No spurious correlations (−0.5 < r < 0.5) were detected between parameters (**Fig. 3b**), allowing for independent and sensitive recovery of each of the seven parameters ^58^. Thus, our model recovered each of the seven parameters independently and sensitively.

**Fig. 3:**
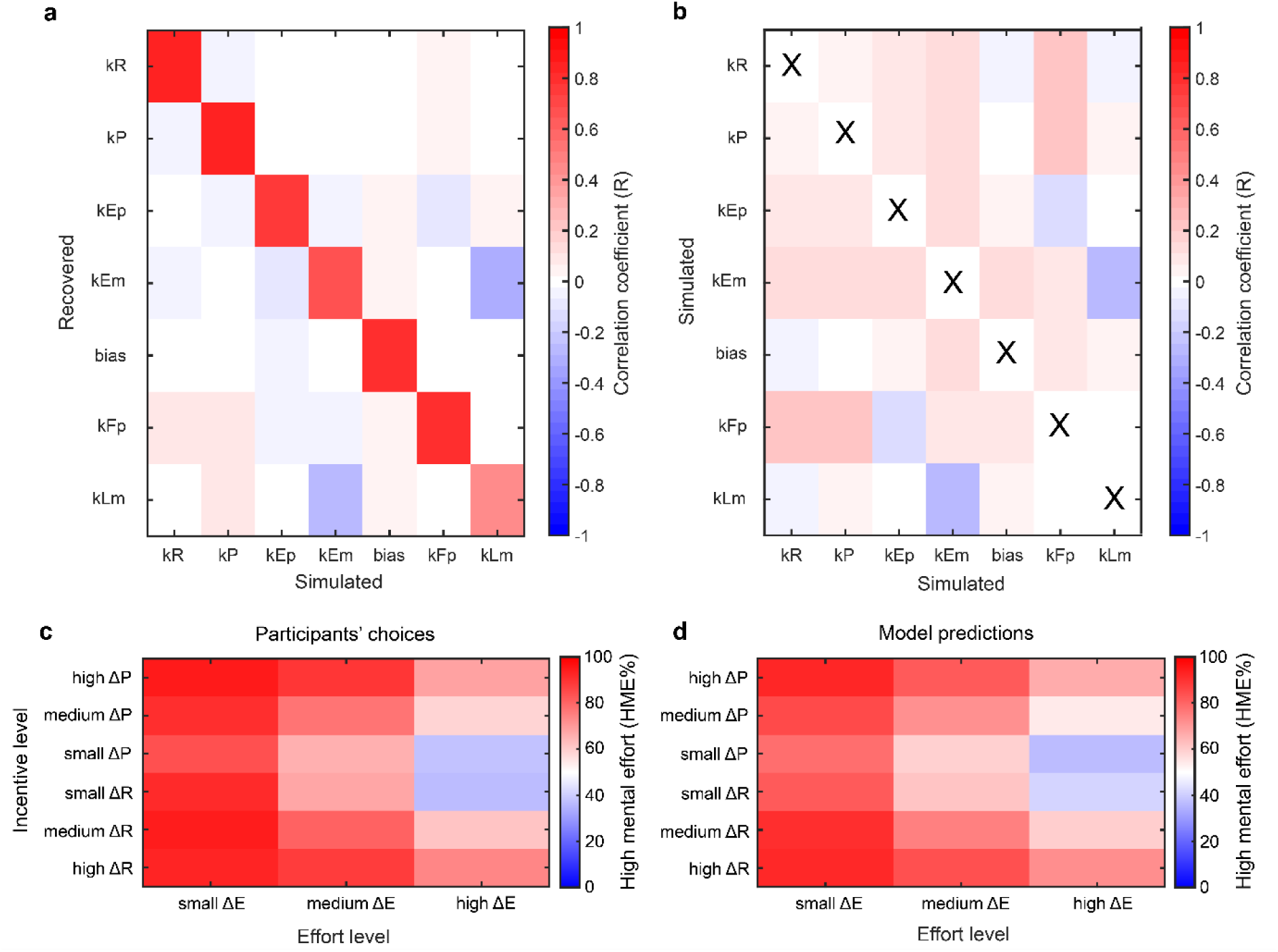
Comprehensive analysis of model parameters. (**a**) Confusion matrix evaluating the model’s parameter recovery capability, contrasting recovered parameters against their original (simulated) counterparts. The parameters correspond to the sensitivity for reward (kR), punishment (kP), physical effort (kEp), mental effort (kEm), a bias toward the high effort option (bias), physical fatigue (kFp) and mental learning (kLm). (**b**) Autocorrelation matrix derived from 30’000 simulations assessing the identifiability of all model parameters. (**c**) Heatmap representing the percentage of high mental effort (HME) choices in function of the difference between the high and the low effort option in terms of effort and incentive levels across participants. (**d**) Corresponding heatmap representing the percentage of high mental effort (HME) choices predicted by our computational model in function of the difference between the high and the low effort option in terms of effort and incentive levels.

Further validation was performed by challenging our behavioral model against variants with fewer parameters (**Fig. S4**). Model comparisons unanimously highlighted the importance of all the extracted parameters (i.e., kR, kP, kEm, kEp, kLm, kFp, bias) in describing participants’ behavior, despite the penalization for the added complexity of each additional parameter in the comparison process. Importantly, our model closely mirrored participant choices (median absolute error of 20.03 ± 7%, goodness-of-fit R^2^ = 0.46) (**Fig. 3c-d**), effectively capturing the tendency of participants to choose more effortful options when monetary incentives were higher or the associated effort was less demanding.

Interestingly, we did not observe any statistical difference across individuals between the reward (kR) and the punishment (kP) sensitivities (Mann-Whitney U test: p = 0.62) suggesting that, at least in our task, effort investment was not more important than loss aversion.

### dmPFC/dACC metabolites specifically explain sensitivity to mental effort

Next, we sought to determine whether dmPFC/dACC metabolites could explain any of the five idiosyncratic parameters characterizing HME modeling from the computational modeling above. Given that cognitive effort is often perceived as aversive, which influences action avoidance ^59^, we initially focused on predicting the sensitivity to mental effort (kEm) from brain metabolites concentrations. The dataset was randomly split into 75% for training/validation (N=53) and 25% for testing (N = 18), with hyperparameter tuning conducted using *hyperopt* and CVLOO. The XGBoost regression model showed a good level of accuracy in predicting kEm (range = 2.13, train RMSE = 0.34, validation RMSE = 0.75). The test set yielded a modest prediction error (test RMSE = 0.64, r = 0.4, p = 0.046) (**Fig. 4a**), accounting for 16% of kEm’s variance. This was confirmed by a permutation test, which indicated predictions significantly above chance level (95^th^ percentile = 0.73 > model RMSE = 0.64) (**Fig. 4b**). In contrast, training a model to predict other parameters involved in mental-effort decision making such as reward sensitivity (kR), punishment sensitivity (kP), mental learning progress (kLm), and bias resulted in no significant predictions (all p > 0.4). This further emphasizes that our model strongly predicts cognitive effort-related behavioral components with both HME and kEm, while it does not succeed to significantly predict other predictors of motivated behavior from our computational model.

**Fig. 4:**
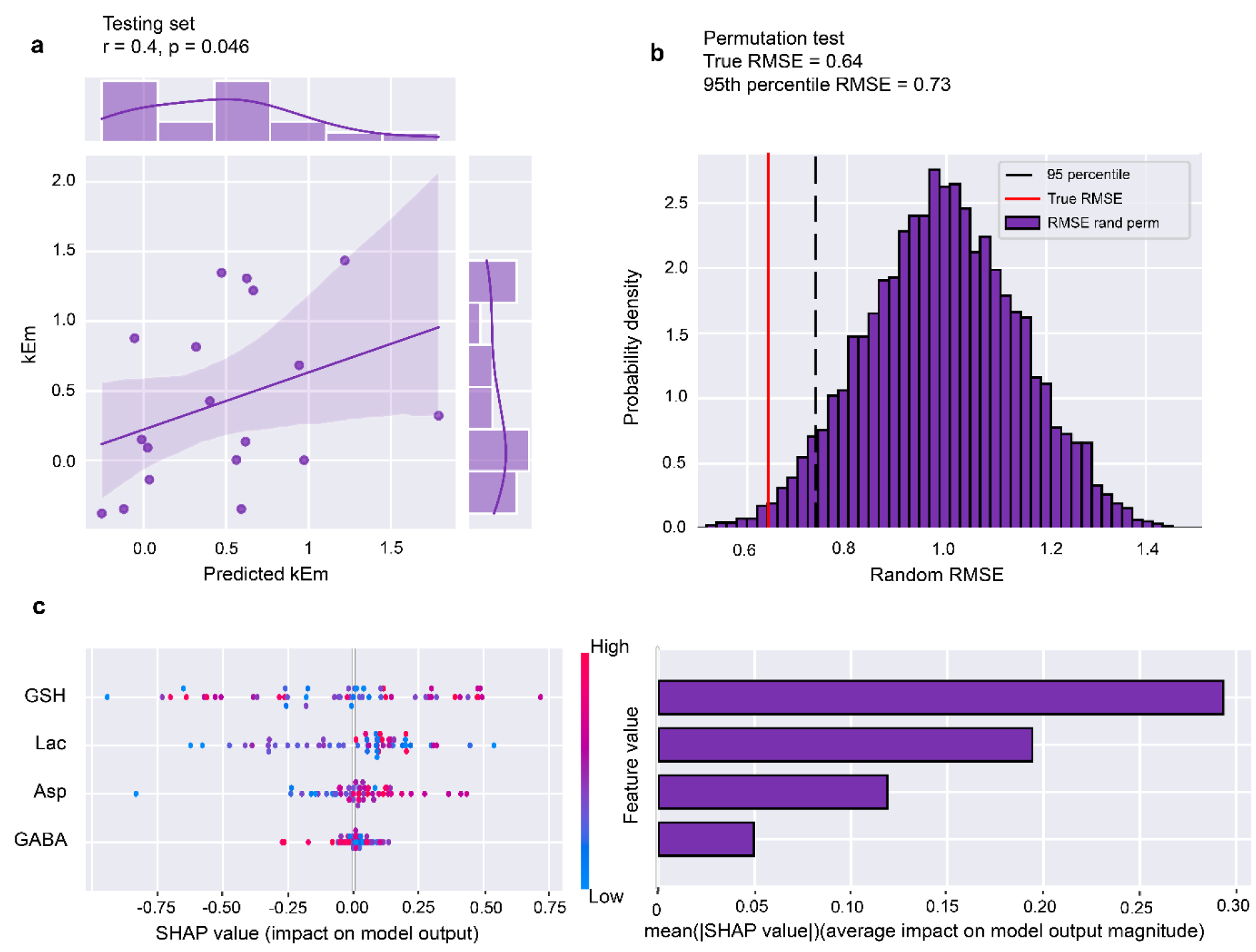
Assessment of the model’s explanatory power for mental effort sensitivity (kEm) using dmPFC/dACC metabolite concentrations. (**a**) A model based on dmPFC/dACC metabolites (model 3) selected through an automated feature selection process shows a positive correlation (r = 0.40, p = 0.046) between predicted and observed values of kEm based on the XGBoost model’s test set performance, highlighting the model’s predictive accuracy. (**b**) Histogram of root mean square error (RMSE) from a permutation test involving random label permutation showing a clear validation of the model’s robustness. The test and the results reveal that the model’s RMSE (0.64) surpasses the 95^th^ percentile of the permuted data (threshold RMSE = 0.73), confirming that the model’s predictive success is not due to chance. (**c**) SHAP summary plot representing the impact of each feature on the model output for individual predictions. The analysis identifies five key metabolite parameters— Glutathione (GSH), Lactate (Lac), Aspartate (Asp) and γ-Aminobutyric acid (GABA). Right panel: The horizontal dispersion of SHAP values per feature indicates the variability and influence of each on model predictions.

SHAP values on the impact of the four metabolites predicting kEm prediction highlighted glutathione (GSH), lactate and aspartate as the most discriminating features (**Fig. 4c**). Whereas there was no significant correlation between kEm and dmPFC/dACC GSH or lactate concentrations (p > 0.05) as can also be observed in the SHAP values (**Fig. 4c**), dmPFC/dACC aspartate concentrations exhibited a positive linear correlation with kEm (r = 0.42, p < 0.001; **Fig. 4c** & **Fig. S5b**).

### Refining the model explaining mental effort decision-making with essential biomarkers

Thus, we have found two effective XGBoost models for HME and kEm (model 2 and 3), utilizing 8 and 4 dmPFC/dACC metabolites, respectively. We next applied a parsimonious approach to determine if a smaller subset of metabolites could predict HME (model 4), aiming to balance simplicity with explanatory power ^60^. This approach aligns with cognitive neuroscience and machine learning practices, where effective models prioritize high predictive accuracy with minimal parameters ^61^. However, selecting the top two (i.e., glutamate and lactate; **Fig. S6a-c**) or top three (i.e., glutamate, lactate, and glutamine; **Fig. S6d-f**) predictors from our SHAP analysis failed to yield a significant prediction.

Inspired by efforts to integrate biological context in machine learning for improved relevance and interpretability ^62^, we examined metabolic overlap among the top-ranked metabolites. While aspartate ranked highly in the kEm model (**Fig. 4c**), its rank was lower in the HME model (**Fig. 2e**), possibly due to its biosynthetic relationship with glutamate and glutamine in the TCA cycle (**Fig. 5**) ^63^. Machine learning’s tendency to minimize correlated features might have undervalued aspartate’s role. Significant intercorrelations among these metabolites (**Fig. S5g**) support this hypothesis, suggesting aspartate may provide unique predictive information despite its lower ranking. In contrast, lactate, uncorrelated with glutamate (r = 0.03, p = 0.77) or aspartate (r = 0.19, p = 0.11), retained distinct variance. This prompted re-evaluation of glutamate, aspartate, and lactate concentrations in the dmPFC/dACC for predicting HME and their role as biomarkers for mental effort decision-making.

**Fig. 5:**
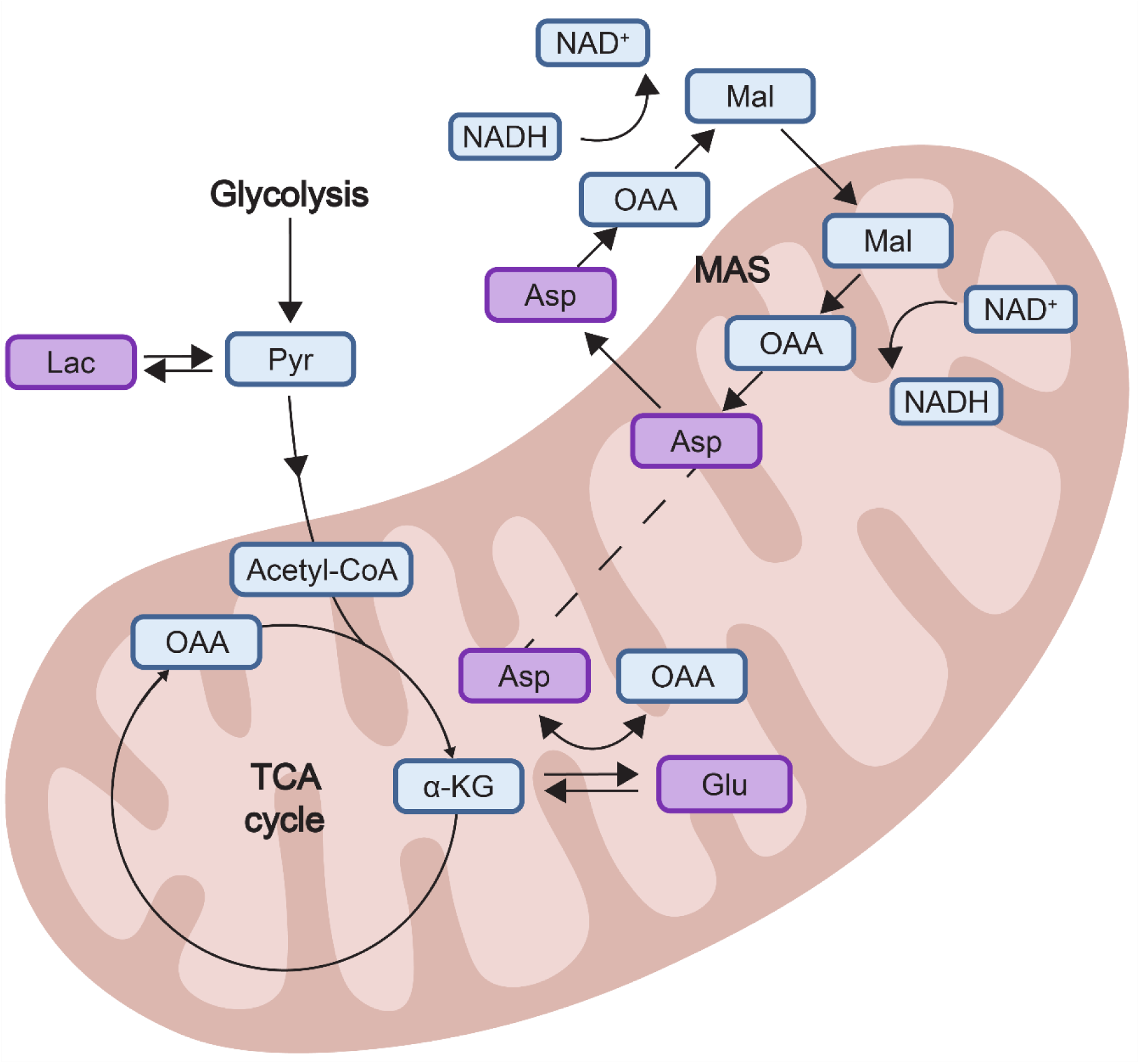
Schematic representation of the identified metabolites within the tricarboxylic acid (TCA) cycle and the malate-aspartate shuttle (MAS) within mitochondria. The scheme depicts the interplay among key metabolites pertinent to neuronal energy metabolism. Highlighted in bold purple are the key metabolites related to mental effort motivation identified in this study: glutamate (Glu), aspartate (Asp), and lactate (Lac). The diagram traces the pathways of Asp and Glu within the TCA cycle and the role of Lac in glycolysis. Key intermediates include pyruvate (Pyr), alpha-ketoglutarate (α-KG), oxaloacetate (OAA), malate (Mal), and acetyl-CoA, along with the redox couple NAD+/NADH, integral to both the TCA cycle and MAS. This visual abstract underscores the metabolic fluxes that are hypothesized to underpin the cognitive processes.

We retrained an XGBoost model (model 4) using a train/validation/test analysis design with a CVLOO approach. Data were randomly split into training/validation (80%, N = 55) and testing (20%, N = 14) sets. XGBoost with RMSE as the error metric, achieved a good fit on the training (RMSE = 0.75%) and validation (RMSE = 15.68%) sets (**Fig. 6a,b**) and performed well on the test set (RMSE = 12.56%, r = 0.64, p = 0.006), explaining 41% of the variance (**Fig. 6c**). A permutation test (5000 permutations) confirmed predictions significantly exceeded chance levels (95^th^ percentile = 13.59 > model RMSE = 12.56). These results indicate that a machine learning model based on a few dmPFC/dACC metabolite concentrations can predict participants’ willingness to exert mental effort.

**Fig. 6:**
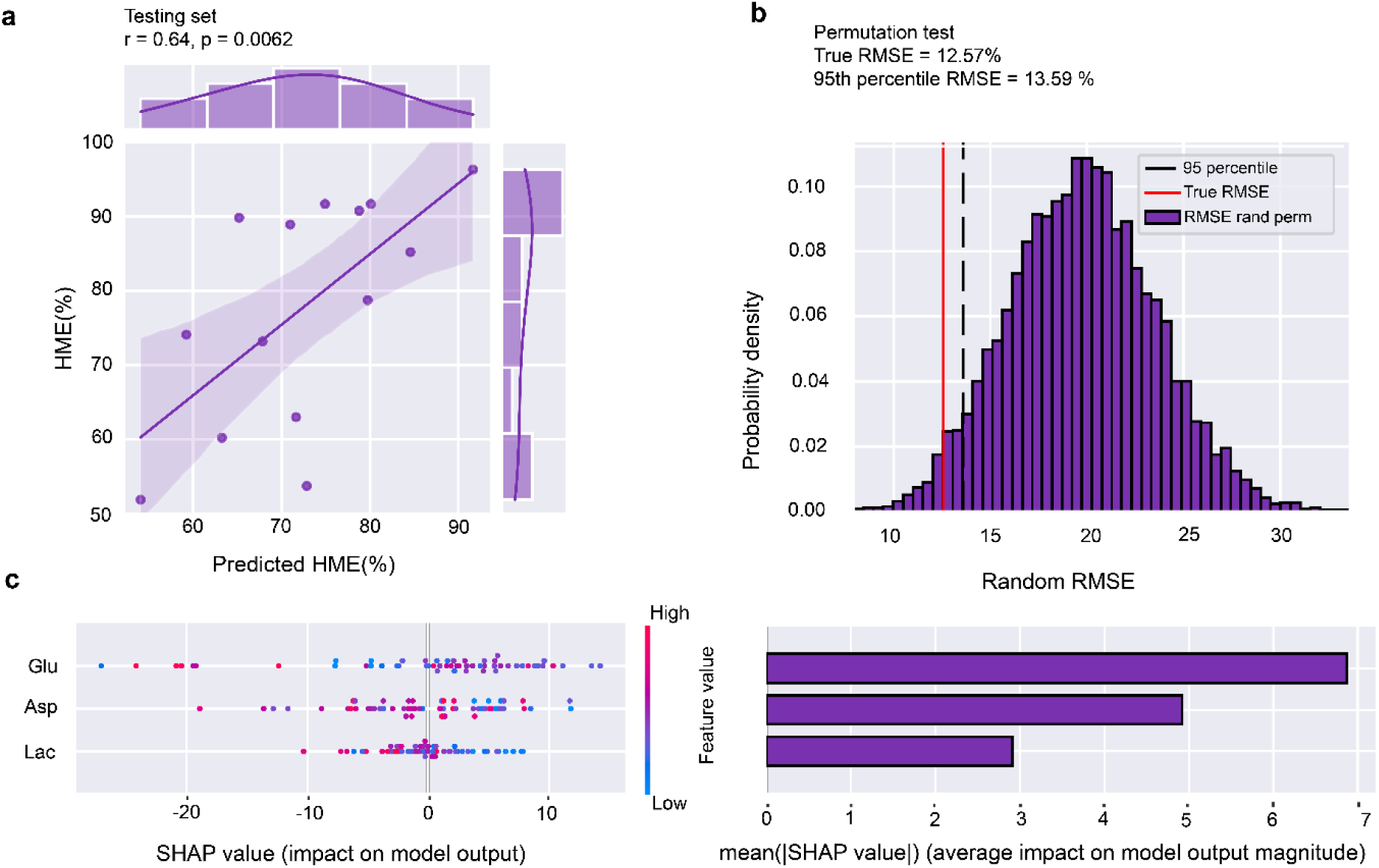
Model for high mental effort (HME) choices using key metabolites from the dmPFC/dACC. (**a**) A strong positive correlation (r = 0.64, p = 0.006) between the XGBoost model’s predicted probability of HME choices (model 4) and the observed percentages within the test set, indicating robust predictive validity using the metabolites glutamate (Glu), aspartate (Asp), and lactate (Lac). (**b**) A permutation test that randomizes the labels and recalculates the RMSE to compare against the model’s true performance indicates the model’s predictive reliability, i.e., the true RMSE of the model (12.57.%) is well below the 95^th^ percentile of the permuted distribution (threshold RMSE = 13.59%), indicating that the model’s predictions are not due to random chance. (**c**) SHAP value summary plot depicting the impact of each metabolite on the model’s predictions for individual subjects. Right panel: The magnitude and distribution of the SHAP values for glutamate, aspartate, and lactate indicate their respective contributions to the model’s decision-making process.

SHAP analysis revealed complex relationships. Glutamate and lactate showed no linear correlations with HME (glutamate: r = -0.19, p = 0.12; lactate: r = -0.13, p = 0.27), though extreme glutamate values suggested a quadratic relationship (**Fig. S3**). Aspartate displayed a strong negative linear correlation with HME (r = -0.37, p = 0.002; **Fig. S5a**) and a strong negative linear correlation with cognitive efficiency defined as the ratio between the number of correct answers (accuracy) and the time it took to solve the 2-back task (speed) during cognitive performance (r = -0.33, p = 0.006; **Fig. S5e**).

### Plasma and brain metabolite concentrations

To investigate relationships between plasma and brain metabolite concentrations, we examined correlations between plasma levels of glutamate, aspartate, and lactate and their counterparts in the dmPFC/dACC and AI (**Fig. S7**). Given glutamate’s lower plasma concentration than its precursor glutamine ^64^, we also analyzed the relationship between plasma and brain glutamine concentrations. No significant correlations were found between plasma and dmPFC/dACC or AI concentrations of glutamate or of aspartate (**Fig. S7a-b**). Lactate showed region-specific correlations (**Fig. S7c**), being significant in the dmPFC/dACC (r = 0.27, p = 0.023), but not in the AI (r = 0.14, p = 0.36) as already reported in our previous publication^36^. Conversely, plasma and brain glutamine levels exhibited a strong positive association in both regions (dmPFC/dACC: r = 0.54, p < 0.001; AI: r = 0.34, p = 0.014; **Fig. S7d**), suggesting a broader brain-plasma relationship.

## Discussion

Differences in individuals’ willingness to undertake incentivized effortful tasks can profoundly influence their life trajectories ^3–5,65^. The dmPFC/dACC and the AI, both part of the salience network ^66–69^, have both been implicated in effort-based decision-making ^2,8,9,11,12,14,70^, but the specific neurobiological factors driving individual variability in this domain remain unclear. Our study bridges this knowledge gap by identifying key neurometabolic predictors in the dmPFC/dACC that account for variability in mental effort exertion.

Departing from earlier studies that focused primarily on individual metabolites ^32,34,39,71–73^, our multivariable approach identified a broader view of the neurometabolic processes associated with mental effort-based decision-making. Using high-field (7T) ^1^H-MRS and machine learning, our initial models identified several metabolites in the dmPFC/dACC as relevant to predicting both high mental effort exertion (HME) and sensitivity to mental effort (kEm). Subsequent refinement of these models revealed that a smaller subset of key metabolites (glutamate, lactate, and aspartate) in the dmPFC/dACC emerged as the most robust predictors of mental effort motivation. Specifically, aspartate and lactate levels correlated positively with aversion to mental effort, while glutamate exhibited an inverted-U shape relationship with HME, suggesting that it may boost motivated behavior within a specific and moderate concentration range. These results also emphasize the complex, nonlinear interactions between neurometabolic markers and mental effort motivation, with the refined model explaining up to 40% of HME variance. Importantly, models using AI metabolites or predicting HPE were inconclusive, supporting a particularly predictive value of dmPFC/dACC metabolites in mental effort-based decision-making in our experiment.

Glutamate, aspartate, and lactate are central to brain metabolism, playing crucial roles in supporting physiological functions critical for cognitive processes ^40,42,74^. These metabolites, interconnected through the tricarboxylic acid (TCA) cycle and glycolysis, underpin neuronal energy supply, neurotransmission, and amino acid synthesis ^75,78,79^. Their combined predictive role for HME suggests that their interplay within the dmPFC/dACC may fine-tune neural computations related to mental effort feasibility and desirability. In addition, the SHAP analysis also identified the importance of other metabolites, such as glutathione related to antioxidant pathways, and glutamine, to a lesser extent. Previous research showed a positive involvement of glutathione in the nucleus accumbens on the regulation of physical effort exertion ^35^, while glutamine levels in the nucleus accumbens were negatively related to physical effort perception ^32^. These findings suggest that metabolic pathways impacting motivated behavior may vary across different brain areas.

Specifically, the neurometabolic profile in the dmPFC/dACC predictive of low motivation for mental effort, involving combinations of elevated levels of glutamate, aspartate, and lactate, suggests a metabolic imbalance potentially indicative of reduced mitochondrial function in some of our participants ^80–82^. Such a disbalance could signal a disruption in the TCA cycle, as suggested by elevated aspartate and glutamate levels ^83^, and a compensatory shift towards glycolysis due to impaired oxidative phosphorylation, indicated by increased lactate levels. These disturbances may impair ATP production, amino acids, nucleotides, and bioactive molecules synthesis, essential for neuronal function and integrity ^26,27^, as well as neurotransmitter synthesis and signaling pathways ^84^. Rodent models suggest that mitochondrial dysfunction impairs performance in demanding behavioral tasks ^26,85,86^, and similar metabolic imbalances within the dmPFC/dACC may affect energy-intensive cognitive processes, diminishing both motivation and cognitive control mechanisms. Our current results shed light on the debate regarding whether cognitive effort relies or not on a biological resource ^87–89^ by emphasizing the essential influence of energetic neurometabolism on the motivation to exert cognitive effort. They also underline that the limited biological resource upon which cognitive effort relies may not be one single molecule, as suggested in former models which proposed glucose ^20^, amyloid-β ^90^, glutamate ^34^ or adenosine ^91^ as the main biological limitation to cognitive effort. Instead, our results suggest that cognitive effort may rely on a combination of several metabolites related to the TCA cycle and to mitochondrial energetic functioning more generally. Future research should therefore study this metabolic pathway rather than trying to single out ‘a’ metabolite as the sole driver/inhibitor of cognitive effort.

Our results were mostly focused on healthy human subjects with variable levels of motivation based on the MADRS-S. In the future, it would be relevant to assess whether they can be confirmed in clinical populations with a diagnosed major depressive disorder and to determine whether these populations indeed suffer from mitochondrial dysfunctions that could underlie their pathology. Moreover, we did not collect information regarding the menstrual cycle phase of female participants in the current study, although it could influence both body and brain metabolism. Future studies could further explore whether there is a relationship between dmPFC/dACC glutamate, aspartate and lactate and the menstrual cycle.

Inspecting the weights of the metabolites within the model provides further insights into their potential contributions, particularly through the observed quadratic association of glutamate with mental effort choices and the strong negative linear correlation of aspartate with high mental effort (HME) choices. Consistent with evidence that both low and high glutamate concentrations can impair cellular functions ^92^ and contribute to cognitive deficits ^93,94^, our study highlights that low or high dmPFC/dACC glutamate levels can impair cognitive effort motivation underlying that they have to be tightly regulated to preserve cellular and cognitive functions. Aspartate’s negative association with HME choices and its link to reduced cognitive efficiency during effortful tasks highlight its significant, yet rather underappreciated role in cognitive metabolism and neuronal activity. Lactate’s contribution to HME and mental effort sensitivity (kEm) becomes evident with the multivariable analysis, with elevated levels potentially reducing HME choices through mechanisms that extend beyond its traditional role as an energy substrate ^95^. Brain lactate levels increase following systemic physiological changes, such as intense exercise ^96,97^, and elevated lactate is associated with pathologies like schizophrenia and depression ^98–101^, which include motivational deficits as core symptoms ^65^.

Notably, while metabolite levels in the brain, exhibit remarkable stability over time ^102–104^, indicating distinct neuro-metabolic phenotypes ^105^, they remain susceptible to changes. For example, stress can dynamically alter glutamate levels in the prefrontal cortex ^106,107^, thereby influencing decision-making ^108,109^. This highlights potential intervention targets for conditions characterized by motivational deficits. Our findings further indicate that brain concentrations of glutamate and aspartate are largely independent of their plasma levels, consistent with prior evidence showing minimal contribution of plasma glutamate to brain levels ^64^. In contrast, the observed positive correlation between plasma and dmPFC/dACC lactate concentrations aligns with the unique transport dynamics of lactate across the blood-brain barrier ^110^. While the possibility of impacting brain concentrations of glutamate and aspartate through peripheral interventions appears limited, the plasma-brain lactate correlation suggests that systemic changes could impact brain metabolism and motivated behavior. Accordingly, interventions such as exercise or direct lactate administration may modulate brain lactate levels, influencing motivated behavior and cognitive engagement.

While our study presents significant insights, it also has certain limitations. Although the models developed for HME and kEm yielded significant and robust predictions, they were trained on a relatively small sample size. This results in an increased sensitivity to randomness parameters such as hyperparameters selection or splitting datasets according to a seed. Thus, our results should be interpreted with caution. However, the recruitment and testing of 75 subjects in the demanding 7T experimental studies represent a considerable sample size for this field. Furthermore, all three models result in similar conclusions, passing the permutation test, further backed by statistical analysis. Additionally, the effort requirements in our tasks were based on varying the duration during which effort had to be sustained, while effort intensity was kept constant, and it would be interesting to assess whether the same results apply to tasks where effort varies in intensity instead (e.g., between 1-back and 3-back). Given the exploratory nature of our results, future research should benchmark the predictive capacity of these models and evaluate their generalizability to new datasets and other mental effort paradigms.

In conclusion, our study advances the understanding of neurometabolism in cognitive neuroscience, particularly in relation to motivated behavior. By identifying the combined predictive roles of glutamate, lactate and aspartate in the dmPFC/dACC, we highlight their collective ability to predict mental effort in healthy individuals. This work bridges the gap between metabolic processes and cognitive function and introduces a novel approach to assess the collective impact of multiple metabolites. Our findings provide a foundation for future investigations, including causal analyses in animal models, and position a subset of dmPFC/dACC metabolites as biological biomarkers for mental effort and motivation.

## Author contributions

Conceptualization: AB, NC, CS. Methodology: AB, JPB, JB, BC, NC, MP, CS, LX. Software: AB, NC, LX. Formal Analysis: AB, JPB, BC, NC. Investigation: AB, JPB, BC, NC. Resources: JPB, BC, CS, LX. Writing-original draft: AB, NC, CS. Visualization: AB. Supervision: NC, CS, LX. Project Administration: CS. Funding Acquisition: NC, CS.

## Acknowledgments

We thank the nursing staff at EPFL-Point Santé (Viviane Depuydt-Linder and Chiyama Mathivathanasekaram) and Centre Medical des Arcades (Lausanne), as well as members of the EPFL-LGC laboratory for their support in blood collection and processing. We thank Antonius Wiehler, Alizée Lopez-Persem and Bogdan Draganski for their inputs to the behavioral experimental design, and Olivier Ciclet (Nestlé Research), Adrien Frezal (Nestlé Research), and Irina Monnard (Nestlé Research) for their contributions to the plasma analyses.

