## Supplementary information for "Neurometabolic predictors of mental effort in the frontal cortex"

### Supplementary Methods

**Participants**

A total of 75 healthy right-handed volunteers (40 females) participated in this study, approved by the Cantonal Ethics Committee of Vaud (CER-VD), Switzerland. Recruitment was done through the Université de Lausanne (UNIL) LABEX platform and local advertisements. Fluency in French (minimum B2 level) was a requirement. Exclusion criteria included being outside the 25 - 40 years age range, regular drug or medication use, history of neurological disorders, and MRI contraindications (e.g., pregnancy, claustrophobia, tattoos near the neck, or metallic implants). Informed consent was obtained from all participants before being invited to the study. Pre-visit, online questionnaires were completed, including the Montgomery Asberg Depression Rating Scale – self-rated (MADRS-S), with a cutoff of 4 differentiating non-depressive from mild/high depressive individuals. This ensured a wide behavior range and sufficient inter-individual variability. Four subjects had to be removed due to incomplete (we could not acquire behavior due to technical reasons in 2 subjects) or incongruent behavioral data (2 subjects always picked up the high effort option with high confidence) leaving a final sample of 71 subjects (35 females; mean age = 30.2 ± 3.4 years; mean weight = 67.5 ± 12.1 kg; mean BMI = 23.0 ± 3.1). Because they didn't complete all the trials of the behavioral experiment, we had to remove two additional participants from the 2$^{nd}$ and 4$^{th}$ machine learning models of high mental effort (HME) choices leaving a final sample of 69 subjects (34 females; mean age = 30.5 ± 3.9 years; mean weight = 67.9 ± 12.6 kg; mean BMI = 22.8 ± 3.1) for these models. However, since there were still enough trials to estimate kEm in these subjects, we kept the sample at N = 71 for the machine learning model 3 in order to maximize the number of subjects used for the estimation of model 3. The results presented in this study are part of a broader study investigating inter-individual differences in motivated behavior, from which a subset of the data has been published [49].

Compensation included a base payment of 70 CHF, an additional 10 CHF per experiment hour, and a fixed amount of 4 CHF for each time they performed a physical or mental maximal performance. To ensure that participants performed their maximal performance, they were told that final payment would rely on their performance on these stages. Maximal performance was measured during calibration (1 physical and 1 mental calibration) and before and after each fMRI session for the corresponding effort type to keep track of eventual changes of maximal performance with fatigue so that participants received 40 CHF in total for these 10 maximal performances. Indifference point measurements and task performance also contributed to earnings, averaging 204 ± 17.4 CHF per participant.

**Experimental procedure**

Task training occurred outside the scanner, with participants' maximum voluntary contraction (MVC) and maximum number of correct responses (MNCR) first determined to calibrate the 4 levels of physical and the 4 levels of mental effort (E0/E1/E2/E3) used in the task. The range of incentives used (P0/P1/P2/P3; R0/R1/R2/R3) was calibrated by identifying one indifference point for each effort type, corresponding to a 50% chance of accepting the level of effort (E2) for a given monetary incentive.

The behavioral task explores different aspects of motivated behavior while capturing a wide range of interindividual differences (**Fig. 1d**). On every trial, participants were asked to choose between a fixed low incentive/low effort option and a high incentive/high effort option, varying in both effort and incentive levels. After their choice selection, the selected option was displayed by continuous (for high confidence) or dotted (for low confidence) lines reflecting the level of confidence indicated by the subject. After their choice, participants had to perform the corresponding effort chosen. Incentives were either framed as monetary loss or gain. Effort was either physical, implying to exert a force equal or superior to 55 % of the participant's MVC during a varying amount of time depending on the level of effort selected, or mental, implying to perform different numbers of correct responses in a 2-back task, depending on their MNCR.

**Physical Training and Calibration**

Physical effort was measured using a handheld dynamometer (TSD121B-MRI, BIOPAC). To determine the maximal voluntary contraction (MVC), participants were instructed to squeeze the dynamometer as hard as possible three consecutive times. The highest value from these 3 attempts was defined as the MVC. Physical effort difficulty fluctuated by adjusting the time interval (0.5 sec for the fixed low effort option E0, and E1: 1.5 sec, E2: 3 sec, and E3: 4.5 sec for the other levels) during which participants had to maintain the force above a predetermined threshold set at 55% of each participant's MVC. To get well acquainted with task difficulty prior to performing the actual task, participants were asked to perform each level of effort five times during the training period, for a total of 20 trials. Additionally, 4 choice test trials followed, to ensure participants understood the task, in particular that each choice would have to be implemented so that they can observe how effortful each level of effort is before engaging with the main task.

**Mental Training and Calibration**

Mental effort was operationalized as a 2-back task. Participants were instructed to answer whether the number displayed on the screen was higher or lower than 5, relative to the number presented two digits earlier. To ensure consistent task performance, participants were first trained to achieve a minimum of 6 correct responses under 10 seconds, with at least 80% accuracy. After a training period lasting between 30 to 185 trials (mean = 66 ± 40), participants were required to achieve the highest possible score in under 10 seconds, which served as a calibration for the difficulty levels. The maximum number of correct responses (MNCR) was defined as 80% of the averaged best score from three attempts, due to the high variability in this measure. To standardize difficulty levels across participants, cognitive effort levels were computed as $\frac{MNCR}{9}$ for the fixed effort level (E0), and $\frac{MNCR}{3}$ (E1), $\frac{2 \cdot MNCR}{3}$ (E2), and MNCR (E3) for the 3 other levels, rounded up to the closest integer. To ensure familiarity with all effort levels, participants performed all levels 5 times (20 total) and completed 4 choice test trials.

**Indifference Point Measurement**

To ensure a widespread response to the task, the incentive levels in the task were adjusted based on the participants' indifference point (IP) determined at the end of the training session, following Westbrook and colleagues [1]. The indifference point was defined as a 50% chance of accepting the level of effort difficulty (E2) for a given monetary incentive [1]. On the first trial, subjects were presented with a low reward (0.5 CHF)/low effort (effort level 0) option and a high reward (1 CHF)/high effort (effort level 2) option. The low reward/low effort option and the effort associated to the high reward/high effort option were constant across the whole procedure, while the reward associated to the high reward/high effort option varied according to the previous choice. The update of the high reward value was done following:

$$R(t+1) \;=\; \begin{cases} R(t) + \dfrac{R(t) - 0.5}{2} \; \textit{when the low reward was chosen in trial } t \\ R(t) - \dfrac{R(t) - 0.5}{2} \; \textit{when the high reward was chosen in trial } t \end{cases}$$

The value of R(t+1) was also bounded at 1 so that the difference would never go above 0.5 between the two options. The indifference point was taken as the value of R(t+1) after 5 trials. This procedure was done independently for both the cognitive and the motor domain. For the sake of time, we only calibrated the indifference point in the gain domain and then used equivalent values in the loss domain.

The range of incentives that we used in the main task was defined as such for each task: $R0 = 0.5\,CHF;\ R1 = IP - \frac{dIP}{2};\ R2 = IP;\ R3 = IP + \frac{dIP}{2};\ P0 = -0.5\,CHF;\ P1 = -0.5 + \frac{dIP}{2}\ ;\ P2 = -0.5 + dIP\ ;\ P3 = -0.5 + \frac{3 \cdot dIP}{2}$ with $dIP = IP - 0.5$.

**The Behavioral Task**

The task comprised four alternating and counterbalanced blocks of physical and mental effort, each consisting of 54 trials, with a total of 216 trials. At the beginning and end of each block, participants completed a maximal capacity assessment of either the MVC or the MNCR (depending on the block type). During each trial, participants underwent two phases: the choice phase and the effort phase.

The choice phase began with a white fixation cross with a jitter (0.5-3.5 seconds), followed by the presentation of two options, a low incentive/low effort option and a high incentive/high effort option that varied in effort and incentive level. The effort levels were represented by a yellow pie chart, and the size of the slice indicated the corresponding effort level (0-3). The position of the low incentive/low effort and of the high incentive/high effort options on the left or right of the screen was counterbalanced across trials. Participants used a four-button response pad to select which option they wanted to choose (low incentive/low effort or high incentive/high effort) between the one presented on the left (two left buttons on the response pad) or the one on the right (two right buttons on the response pad). Participants were also requested to provide their confidence in this choice by selecting either the middle buttons to indicate low confidence or the buttons at the extremes of the response pad to indicate high confidence in the choice (**Fig. 1d**). If participants did not make a choice, the low incentive/low effort option was selected by default, and that trial was discarded from the analysis. After the choice selection, the selected option and confidence were displayed for 2 seconds.

The effort phase began with a black fixation cross with a jitter (0.5-3.5 seconds), followed by a physical effort or mental effort task, during which participants had to produce a fixed level of force based on their MVC or solve a certain number of 2-back stimuli, respectively. For the physical effort task, a vertical bar representing the participant's real-time force was displayed on the left, overlaid with a red limit representing the required threshold to exceed (corresponding to the participant’s 55% of MVC). The amount of effort left was displayed as a slice of the pie chart in the center of the screen. If participants exceeded the threshold, the yellow circle diminished over time until it disappeared, ending the effort phase. If not, the yellow pie chart froze, waiting for participants to exceed the threshold again. For the mental effort task, the same pie chart was displayed, along with the current number of 2-back stimuli. Each correct response removed a portion of the pie chart, while incorrect responses increased the number of correct responses required, preventing participants from answering randomly. At the end of the effort phase, the screen displayed feedback on the amount of money won (or lost) on the current trial.

To further reduce the influence of risk on effort discounting (i.e. that participants don't choose a certain option by fear of failure rather than because of effort aversion), even when participants did not achieve the requested effort within the allotted time, participants still obtained a proportion of the reward at stake (or avoided a proportion of the punishment at stake) according to their percentage of effort completion achieved.

**Computational modeling of motivated behavior**

Participants' choices were fitted with a softmax model using Matlab's VBA toolbox (https://mbb-team.github.io/VBA-toolbox/) which implements Variational Bayesian analysis under the Laplace approximation [2]. The algorithm provides an estimate of the posterior density over the model parameters, starting from Gaussian priors.

We developed four models to fit participants' choices in our task. We designed these models to increase in complexity and then, based on model comparison, be able to select which model was best to explain participants' behavior. Confidence was included in the choice outcome (0=low effort chosen with high confidence; 0.25 = low effort chosen with low confidence; 0.75 = high effort chosen with low confidence; 1 = high effort chosen with high confidence) to improve the accuracy of our computational models. To reduce the risk of overfitting, and improve robustness, all models' components were kept linear. They all aimed at fitting the probability of choosing the high incentive/high effort option ($P_{High\ effort}$) based on a *softmax* transformation of a given difference in subjective value (ΔSV) for a given trial *t* where ΔSV is the subjective value of the high incentive/high effort option discounted by the subjective value of the low incentive/low effort option. We started with a basic model that only considered incentive values (model 1). Our second model took both incentive and effort into account (model 2). Following another publication [3], we also modeled a general tendency in performing the task independent of the task variables with a *bias* constant in our third model (model 3). Our fourth model considered the effect of effort over time which could either promote (learning) or decrease (fatigue) the subjective costs of selecting the costly option. The details for each model were as follows:

**Model 1:** kIncentive (kI) model predicting that choice only depends on the monetary incentives.

$$\Delta\mathrm{SV(t)} = \ \mathrm{kI} \cdot \Delta\mathrm{I(t)}\ (\mathrm{Eq.\,1})$$

$$\mathrm{kI} * \Delta\mathrm{I(t)} = \mathrm{kR} \cdot \Delta\mathrm{R(t)} + \mathrm{kP} \cdot \Delta\mathrm{P(t)}\ (\mathrm{Eq.\,2})$$

where ΔR and ΔP represent the difference in the monetary incentives (reward or punishment) between the varying option and the fixed option (ranging from 0.01CHF to 0.49 CHF for ΔR, and -0.01 to -0.49CHF for ΔP), kR represents the sensitivity to rewards and kP the sensitivity to punishments.

$$\mathrm{Pr}_{\mathrm{High\ effort\ choice}}\ (\mathrm{t}) = \frac{1}{1 + \mathrm{e}^{-\Delta\mathrm{SV(t)}}}\ (Eq.3)$$

**Model 2:** kI + kEffort (kE) model predicting that choice depends on both the monetary incentive and the effort levels.

$$\Delta\mathrm{SV(t)} = \ \mathrm{kI} \cdot \Delta\mathrm{I(t)} - \mathrm{kE} \cdot \Delta\mathrm{E(t)}\ (\mathrm{Eq.}4)$$
$$\mathrm{kE} * \Delta\mathrm{E(t)} = \mathrm{kEp} \cdot \Delta\mathrm{Ep(t)} + \mathrm{kEm} \cdot \Delta\mathrm{Em(t)}\ (\mathrm{Eq.}5)$$

Where kEp is the sensitivity to physical effort, kEm the sensitivity to mental effort, ΔEp the difference between the physical effort level of the high incentive/high effort option (1, 2 or 3) and the fixed low effort level (0) and ΔEm the difference between the mental effort level of the high incentive/high effort option (1, 2 or 3) and the fixed low effort level (0).

$$\mathrm{Pr}_{\mathrm{High\ effort\ choice}}\ (\mathrm{t}) = \frac{1}{1 + \mathrm{e}^{-\Delta\mathrm{SV(t)}}}\ (Eq.6)$$

**Model 3:** kI + kE + Bias model predicting that choice depends on monetary incentives, the effort levels and an individual bias towards the high (or the low) effort option to account for a general bias towards performing the task which could relate to intrinsic motivation.

$$\Delta\mathrm{SV(t)} = \ \mathrm{kI} \cdot \Delta\mathrm{I(t)} - \mathrm{kE} \cdot \Delta\mathrm{E(t)}\ (Eq.7)$$
$$\mathrm{Pr}_{\mathrm{High\ effort\ choice}}\ (\mathrm{t}) = \frac{1}{1 + \mathrm{e}^{-\Delta\mathrm{SV(t)+bias}}}\ (Eq.8)$$

Where *bias* is a constant term capturing a general tendency to choose (or to avoid) the high effort option, independently of the monetary incentives and of the effort levels at stake.

**Model 4:** kI + kE + Bias + kTime integrated model predicting that choice depends on monetary incentives, the effort levels, an individual bias towards the high (or the low) effort option and a variable fluctuating with time (possibly reflecting physical fatigue in the physical domain and mental learning progress in the mental domain).

$$\Delta\mathrm{SV(t)} = \ \mathrm{kI} \cdot \Delta\mathrm{I(t)} - \Delta\mathrm{E(t)} \cdot \left(\mathrm{kE} + \mathrm{kF} \cdot \mathrm{T}_{\mathrm{integrated}}\right)\ (Eq.9)$$

$$\mathrm{kF} * \mathrm{T}_{\mathrm{integrated}} = \mathrm{kFp} \cdot \sum_{\mathrm{t=0}}^{\mathrm{t-1}} \mathrm{AUC(t)} - \mathrm{kLm} \cdot \mathrm{Efficiency(t-1)} (Eq.10)$$

Where kFp represents the sensitivity to physical fatigue and $\sum_{t=0}^{T} \mathrm{AUC(t)}$ is the sum of the integral of the force exerted in all the previous trials of the current session as measured by the area under the curve (AUC) of the force exerted during the effort phase for any given trial *t*:

$$\mathrm{AUC(t)} = \int exerted\, force(t)\ (Eq.11)$$

kLm represents the mental learning sensitivity and $Efficiency(t-1)$ represents the mental efficiency in the preceding trial (t-1). Efficiency is defined the ratio between the number of correct answers and the total trial time in the previous trial:

$$\mathrm{Efficiency(t-1)} = \frac{\mathrm{N.}correct\ (t-1)}{\mathrm{Total\ trial\ time(t-1)}} (Eq.12)$$

The ΔSV formula can also be written more extensively as:

$$\Delta SV(t) = kR * \Delta R(t) * idx_I + kP * \Delta P(t) * (1 - idx_I) - (kEp * \Delta Ep(t) + kFp * \Delta Ep(t) * \sum_{t=0}^{T} AUC(t)) * idx_E - \left(kEm * \Delta Em(t) + kLm * \Delta Em(t) * \frac{\mathrm{N.correct}\ (t-1)}{Total\ trial\ time(t-1)}\right) * (1 - idx_E) \quad \text{(Eq. 13)}$$

Where $idx_I$ is an index indicating the incentive condition of the current trial ($idx_I$=1 for rewards and $idx_I$ =0 for punishments) and $idx_E$ is a binary index indicating the effort type of the current block ($idx_E$=1 for physical and $idx_E$=0 for mental blocks).

Finally, ΔSV is then taken as input in a *softmax* function:

$$\mathrm{Pr_{High\ effort\ choice}\ (t)} = \frac{1}{1 + \mathrm{e}^{-\Delta \mathrm{SV(t)} + \mathrm{bias}}}\ (Eq.14)$$

Apart from the bias parameter (*bias*), all parameters were positively constrained through a log(1+exp(x)) transformation. All parameter priors were attributed with a null mean and a sigma of 1, except for kR and kP where, due to high fluctuations in ΔI across participants (depending on IP calibration), we decided to set the sigma to 100. The mean value of the estimated parameters was then used for the subsequent analyses.

We also developed one more model to test whether a parabolic accounting of effort costs would provide a better fit of individuals behavior. The model was identical, except for the effort cost terms (highlighted in bold in the following Equation 15):

$$\Delta SV(t) = kR * \Delta R(t) * idx_I + kP * \Delta P(t) * (1 - idx_I) - (kEp * \boldsymbol{\Delta Ep(t)^2} + kFp * \Delta Ep(t) * \sum_{t=0}^{T} AUC(t)) * idx_E - (kEm * \boldsymbol{\Delta Em(t)^2} + kLm * \Delta Em(t) * \frac{\mathrm{N.correct}\ (t-1)}{Total\ trial\ time(t-1)}) * (1 - idx_E) \quad \text{(Eq. 15)}$$

#### Model comparison

For each model comparison performed with the VBA toolbox [2], we compared the results of the exceedance probability, estimated model frequency, BIC, and AIC, which explicitly penalize the usage of too many parameters.

#### Blood sampling and metabolite acquisition

From our blood sampling, a vial was kept on ice and processed in our laboratory in S-monovette 1.2 mL tubes (Sarstedt, Nümbrecht, Germany) containing EDTA K2E to avoid blood clotting. Whole blood was transferred into a Falcon tube coated with EDTA K2. Subsequently, the tubes were centrifuged at 1100G at 4°C for 15 minutes (2400 rpm in the PK 120R ALC centrifuge). Following centrifugation, 100 µl aliquots of plasma were prepared, to which 5 µl of the protease inhibitor cocktail (PIC) were added.

Plasma lactate concentrations were obtained using an ultrahigh-performance liquid chromatography instrument (Waters Aquity, Milford, MA, USA) coupled to a tandem mass spectrometer (Sciex 6500+, Toronto, Canada). Briefly, 40 µL of plasma samples were deproteinized and then derivatized with 3-nitrophenylhydrazine (3-NPH) according to Dei Cas and colleagues [4]. Derivatized compounds were then quantified by reversed-phase liquid chromatography on a Raptor ARC-18 UHPLC column (2.1 x 100 mm; 1.7 µm) and analytes were detected in negative mode using multiple reaction monitoring (MRM) transitions specific for each analyte. The concentration of lactate was then calculated by comparison of the ratios between the signal intensity of lactate to labeled lactate used as internal standard, via its corresponding calibration curve.

Plasma amino acids (glutamine, glutamate, aspartate) were analyzed using an ultrahigh-performance liquid chromatography instrument (Waters Aquity, UK) coupled to a tandem mass spectrometer a Xevo TQ-XS triple quadrupole mass spectrometer (Waters, Milford, MA, USA). Briefly, plasma samples were thawed at room temperature, vortexed, transferred into a polypropylene plate, and precipitated with a solution containing labeled internal standards in methanol + 0.1% formic acid (FA) before being centrifuged at 2500 rpm for 10 min. Then, the supernatant was collected for the derivatization step in borate buffer at pH 8.8 with Aminoquinolyl-*N*-hydroxysuccinimidyl carbamate (AcQTag) at 55°C for 10 min and agitated at 500 rpm. Finally, samples were diluted 50 times with a 10mM solution of ammonium formate and 0.1% FA before LC-MSMS analysis [5]. Separations were performed on an AccQtag Ultra $C_{18}$ column, 1.7 µm, 2.1 x 100 mm (Waters Milford, MA, USA), and derivatized amino acids were detected using MRM mode. Data were acquired using MassLynx software (Waters, Wilmslow, UK), and chromatographic peaks were integrated with TargetLynx (Waters, Wilmslow, UK). The concentration of each analyte was then calculated by comparison of the ratios between the signal intensity of each analyte to its corresponding labeled analyte used as internal standard, via their corresponding calibration curve.

**MRS acquisition and preprocessing**

Proton magnetic resonance ($^1$H-MR) spectra were collected using a 7 Tesla/68 cm MR scanner (Magnetom, Siemens Medical Solutions, Erlangen, Germany) with a single-channel quadrature transmitter and a 32-channel receive coil (Nova Medical Inc., MA, USA). To optimize the magnetic field homogeneity, shimming was performed using FAST(EST)MAP [6] for both first and second-order. All 1H MR spectra were acquired using Semi-adiabatic spin echo full intensity acquired localized (sSPECIAL) [7,8] with the following parameters: TE/TR = 16/5000ms, 2 averages, VOI = 20 x 20 x 20 mm for dmPFC/dACC and 20 x 20 x 20 mm for AI dmPFC/dACC. All spectra were corrected for frequency/phase shifts and averaged. LCModel was used for spectral fitting and metabolite quantification [9] with a basis set that included simulated metabolite spectra and an experimentally measured macromolecule baseline. An unsuppressed water spectrum was acquired and used as an internal reference for metabolite quantification in LCModel.

T1-weighted images were acquired by MP2RAGE and used to calculate the tissue composition within the MRS voxel. The T1-weigthed images were then segmented into grey matter (GM), white matter (WM), and cerebrospinal fluid (CSF) in SPM12 toolbox (Wellcome Trust Center for NeuroImaging, London, UK) with the MarsBaR package (https://marsbar-toolbox.github.io/) [10], creating the VOI mask for the MRS voxel. Metabolite concentrations were adjusted for the CSF fraction, assuming water concentrations of 43,300 mM in GM, 35,880 mM in the WM, and 55,556 mM in the CSF. In addition, to ensure all voxels were positioned correctly, we computed a density map of metabolites measurement, highlighting the precision of our voxel positioning, based on our MP2RAGE image and VOI mask (**Fig. S1**).

Following metabolite quantification in LCModel and correction, any computed metabolite concentrations with a Cramér-Rao lower bounds (CRLB) higher than 50 % were rejected. In addition, any participants with metabolite concentrations higher than the median ± 3SD were removed as outliers. Then, the metabolite ratios were computed, namely glutamine to glutamate (Gln/Glu) and glutamate to γ-aminobutyric acid (Glu/GABA). Finally, using the Python package sklearn.IterativeImputer, the missing dataset values were filled by multivariate imputation.

**$^1$H-MRS acquisition and voxel positioning**

Participants' head movements were minimized by positioning foam pieces around their heads. The dmPFC/dACC voxel (20 x 20 x 20 $mm^3$) was placed using T1-weighted magnetization prepared 2 rapid gradient echo (MP2RAGE, TE/TR = 1.88/6000 ms, TI1/TI2 = 800/2700 ms, α1/α2 = 7°/5°, slice thickness = 1 mm, FOV = 192 × 192 × 192 $mm^3$, matrix size = 192 × 192 × 192, bandwidth = 240 Hz/Px) by aligning it on the midline of the axial and coronal planes. In the sagittal plane, the voxel horizontal borders were defined to be parallel to the cingulate sulcus. The anterior vertical border was defined to be above the genu of the corpus callosum. The left anterior insula voxel (20 x 20 x 20 $mm^3$) was aligned to the anterior and to the superior peri-insular sulcus[11] on the sagittal plane. On the axial and coronal plane, we moved the voxel to maximize the signal quality and the number of voxels overlapping the insula, while minimizing contact with the ventral pallidum. Semi-adiabatic spin echo full intensity acquired localized (sSPECIAL) was used to acquire 50 MR spectra blocks (2 average/block) per region of interest, with TE/TR=16/5000ms, spectral bandwidth 4000 Hz, and a number of points of 2048. After LCModel spectral analysis, we extracted 13 metabolites, but we focused on the following metabolite of interest and computed their ratio: aspartate (Asp), γ-aminobutyric acid (GABA), creatine + phosphocreatine (Cr+PCr), glutamine (Gln), glutamate (Glu), glutamine + glutamate (Glx), glutathione (GSH), glycine (Gly), myo-inositol (Ins), lactate (Lac), N-acetylaspartate (NAA), glutamine to glutamate ratio (Gln/Glu), glutamate to γ-aminobutyric acid ratio (Glu/GABA).

Participants performed two sessions in the scanner. Session 1: baseline metabolite concentrations were acquired by proton magnetic resonance spectroscopy ($^1$H-MRS), both in the dmPFC/dACC and the AI. Then, participants were trained out of the scanner to perform our behavior task. Session 2: the participant performed the behavioral task during fMRI acquisition.

**Quantification and Statistical Analysis**

Detailed parameters from statistical tests are reported in figure legends and tables indicating the level of significance, and whether it was corrected after multiple comparisons. All correlations were performed using Pearson correlations. Correlations between regressors and the prediction from our machine learning model were one-tailed, using the *scipy* 1.10.0 toolbox on Python 3.10.6. All other correlations were two-tailed, and were analyzed with MATLAB Version: 9.8.0.1323502 (R2020a). All unpaired t-test were two-tailed. The mean and SD for all the parameters of the computational model can be found in **Table S1** after boxcox transformation.

**Hyperparameter optimization process**

*Hyperopt* hyperparameter selection requires testing different ranges of parameters. Parameter ranges are dependent on the dataset and features selected. As these ranges influence the starting point of error reduction for the machine learning (ML) computation, and thus hyperparameter selection, small fluctuations may be necessary to avoid locking in a local minima in optimization plane in *hyperopt*. The following parameters were used for our three main significant models:

| **Parameters model 2 (HME)** | **Range** |
|---|---|
| Learning rate | [0.01-0.3] |
| Max_depth | [5-50] |
| Min_child_weight | [0-0.2] |
| Gamma | [0-0.2] |
| Reg_lambda | [0-0.06] |
| Reg_alpha | [0-0.06] |
| **Parameters model 3 (kEm)** | **Range** |
| Learning rate | [0.01-0.3] |
| Max_depth | [5-50] |
| Min_child_weight | [0-0.1] |
| Gamma | [0-0.1] |
| Reg_lambda | [0-0.37] |
| Reg_alpha | [0-0.5] |
| **Parameters model 4 (HME)** | **Range** |
| Learning rate | [0.01-0.3] |
| Max_depth | [5-50] |
| Min_child_weight | [0-0.2] |
| Gamma | [0-0.81] |
| Reg_lambda | [0-0.1] |
| Reg_alpha | [0-0.1] |

Furthermore, each run of *hyperopt* selection used 10 max evaluations and underwent 100 boosted rounds in the XGBoost cross validation, with 5 early stopping rounds.

### Supplementary Results

**Computational modeling**

To more precisely quantify changes in the valuation of effort and incentive across the task, we developed a computational model of decision-making. We hypothesize that subjective value is dependent on multiple components. To ensure the full capture of behavior from our computational modeling, we compared several models by adding components step by step in increasing model complexity (**Fig. S4**). Our first computational model of behavior only takes incentives into account by only modeling a sensitivity to rewards (kR) and a sensitivity to punishments (kP), with a higher kR (or kP) value meaning that people weigh more monetary rewards (or losses) compared to the others. The second behavioral model also takes both physical and mental effort into account by adding a sensitivity to physical effort (kEp) and a sensitivity to mental effort (kEm), with a higher kEp (or kEm) meaning that participants perceived physical (or mental) efforts as more aversive. The third behavioral model includes a general bias in choices (Kurniawan et al., 2021). The fourth and final behavioral model takes into account participants' exerted effort over time specifically into account by adding one variable for physical fatigue (kFm) and one to take into account the increase of effortful choices in the mental domain which could be related to mental learning (kLm). Participants with a higher sensitivity to physical fatigue (kFp) found that the proposed effort was increasingly aversive and therefore reduce their selection of effortful options over time, while participants with a higher sensitivity to mental learning (kLm) found mental effort less aversive if they performed well at the previous trial. The behavioral model comparison highlights that the full model, considering the effect of the incentive, effort, time, and general bias in choice, is a better predictor of choice than simpler models in terms of both AIC and BIC, but also exceedance probabilities and estimated frequency (**Fig. S4**). This, in turn, shows that each component is necessary to understand participants' choices. Looking at our full model's performance, we also show that our final model has the highest amount of variance explained and the smallest mean absolute error (MAE) (**Fig. S4**). Our results highlight that the most complex model, including all the parameters of interest, was the best and it was therefore selected for further analysis (**Fig. S4**).

Given that parabolic models are sometimes observed as better fitting individuals' behavior in effort-based decision-making tasks [12,13], we also compared our model where effort is modeled linearly to a parabolic model where the effort cost varies quadratically. However, this model comparison yielded a better score for the simple linear model (linear model: AIC = 189.458, BIC = 200.972, R2 = 0.460, estimated frequency = 0.992, exceedance probability = 1, MAE = 0.202; parabolic model: AIC =

196.390, BIC = 207.903, R2 = 0.424, estimated frequency = 0.008, exceedance probability = 0, MAE = 0.209) thereby confirming that this simpler model fitted better the behavior.

**Predicting HME using only the two best features of the model**

Following the results of our first machine learning model aiming at predicting behavior based on brain metabolites, we questioned whether a subset of the most important features was sufficient to train a model robustly predicting HME. Selecting the two best features, namely glutamate and lactate (**Fig. 2e**), did not result in significant predictions (r = 0.23, p = 0.21; **Fig. S6a-c**). Additionally, selecting the three best features by adding glutamine (**Fig. 2e**) also did not yield significant predictions (r = 0.35, p = 0.11; **Fig. S6d-e**). Due to the inter-correlation between metabolites, important metabolites may have been ranked lower by the model and should be considered which is what we decided to do next (**Fig. 6**).

**Metabolites and performance**

Based on our third model's results, we further questioned whether our metabolites of interest, glutamate, aspartate, and lactate would be related to performance measurements during effort exertion in the mental task (**Fig. S5c-f**). Indeed, we found that participants with higher dmPFC/dACC aspartate levels not only displayed lower HME choices (**Fig. S5a**) and a higher kEm (**Fig. S5b**), but they also expressed a lower mental calibration performance (r = -0.34, p = 0.005; **Fig. S5c**) and overall lower efficiency in the mental effort task (r = -0.33, p = 0.005; **Fig. S5e**). Furthermore, participants with higher aspartate concentrations also took more time to make a choice (r = 0.25, p = 0.046; **Fig. S5d**) suggesting that higher levels of dmPFC/dACC aspartate reflect a general decrease in mental effort capacities that is also reflected in decision-making speed. Additionally, higher lactate concentrations correlated positively with the latency to start performing the mental effort (r = 0.25, p = 0.048; **Fig. S5f**), possibly also reflecting a decrease in mental effort capacities. Thus, while the focus of this current study is on decision-making, our metabolites of interest further correlate to the task performance measurements suggesting that dmPFC/dACC metabolites may influence motivation because of their influence on cognitive capacities.

### Supplementary Tables

| Parameter name | kR | kP | kEm | kEp | Bias | kLm | kFp |
|---|---|---|---|---|---|---|---|
| Mean | 0.48 | 0.25 | 3.88 | 1.38 | 1.90 | 3.4 | 0.14 |
| SD | 0.8 | 0.3 | 3.0 | 1.1 | 1.76 | 3.6 | 0.15 |

**Table. S1:** Mean and SD of parameters scores, prior to boxcox transformation.

#### Supplementary Figures

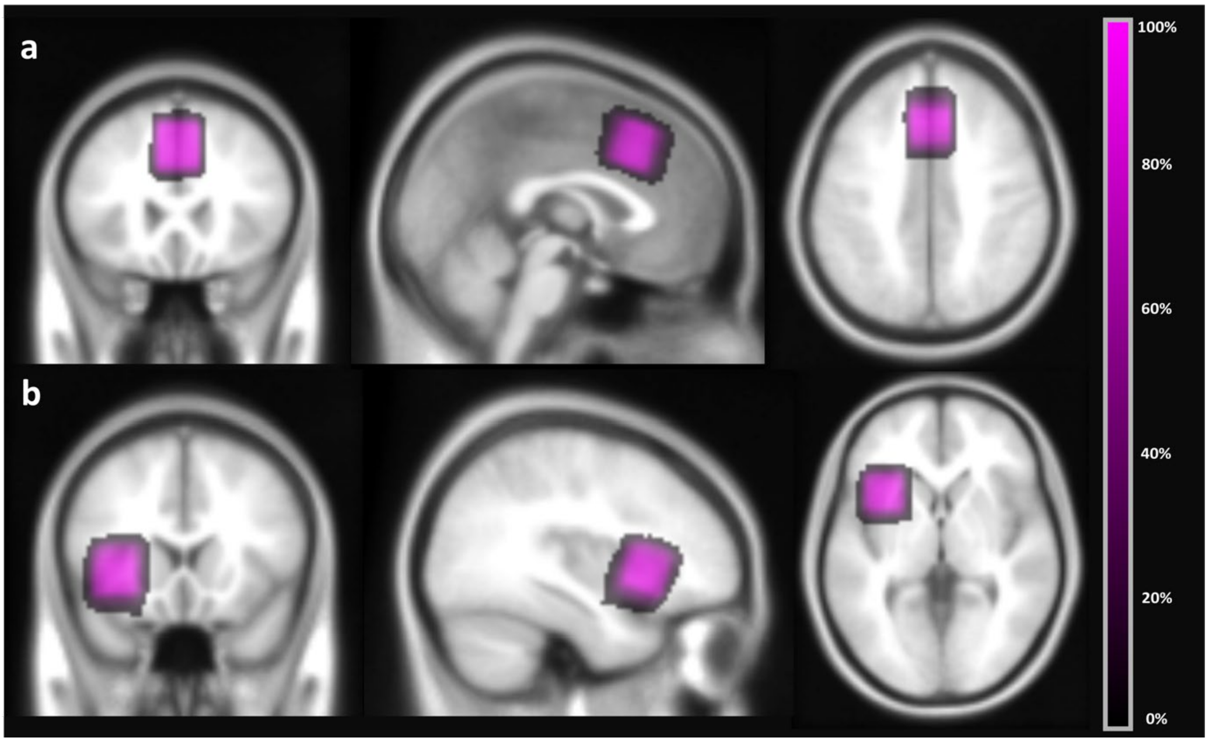


**Fig. S1: $^1$H-MRS voxel density image.** The voxel density maps are overlaid on the MNI152 anatomical template provided in SPM12. (**a**) Voxel density maps in the dmPFC/dACC. (**b**) Voxel density maps in the AI.

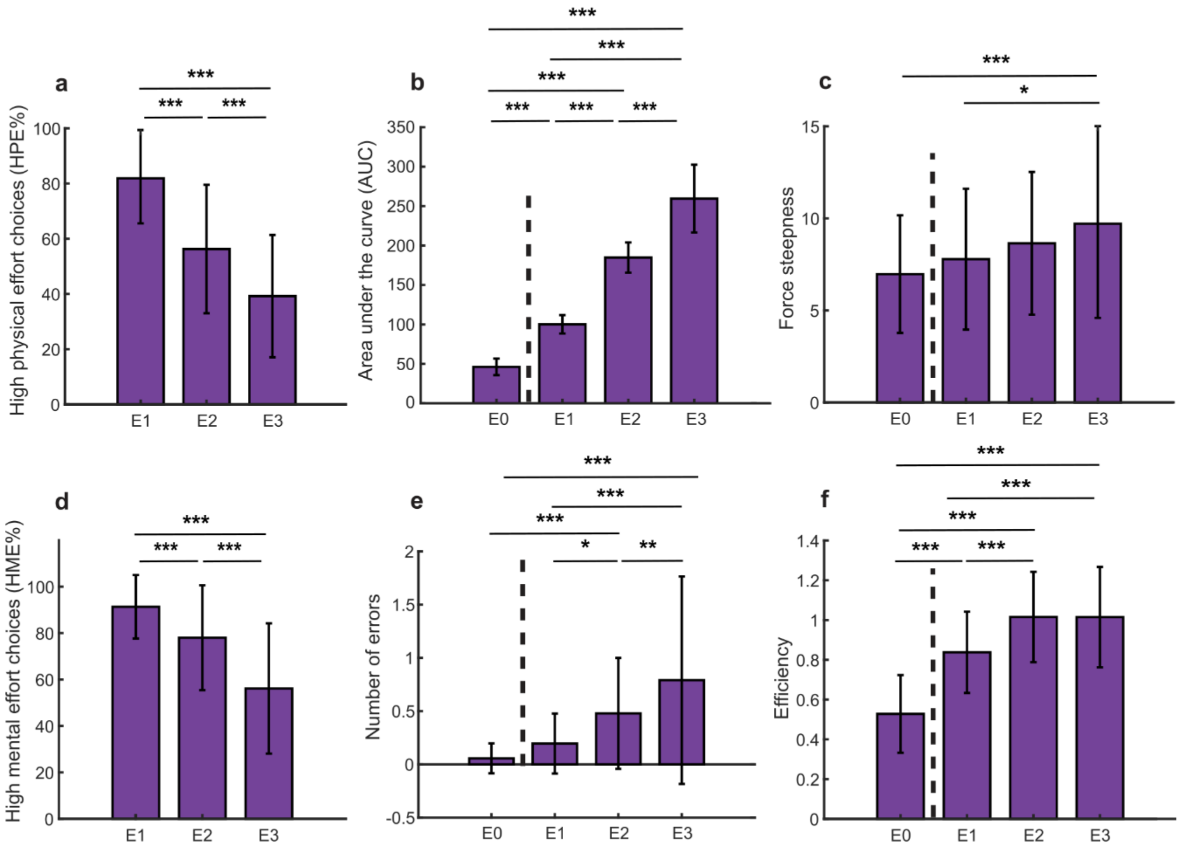


**Fig. S2: Effect of effort level on choice and performance.** In all figures, a dashed line separates the low effort (E0) presented on every trial by default from the high effort option levels (E1-3). (**a)** Percentage of high physical effort choices (HPE), (**b**) area under the curve (AUC) of the produced force and (**c**) initial force steepness increase, are displayed across physical effort levels. (**d**) Percentage of high mental effort choices (HME), (**e**) number of errors, (**f**) and efficiency computed as the number of correct answers divided by the total amount of time to perform a trial, are displayed across mental effort levels. Significance was assessed with a one-way ANOVA followed by a post hoc Kruskal Wallis test. Significance levels: *P < 0.05, **P<0.01, *** P< 0.005, after Bonferroni correction, two-tailed.

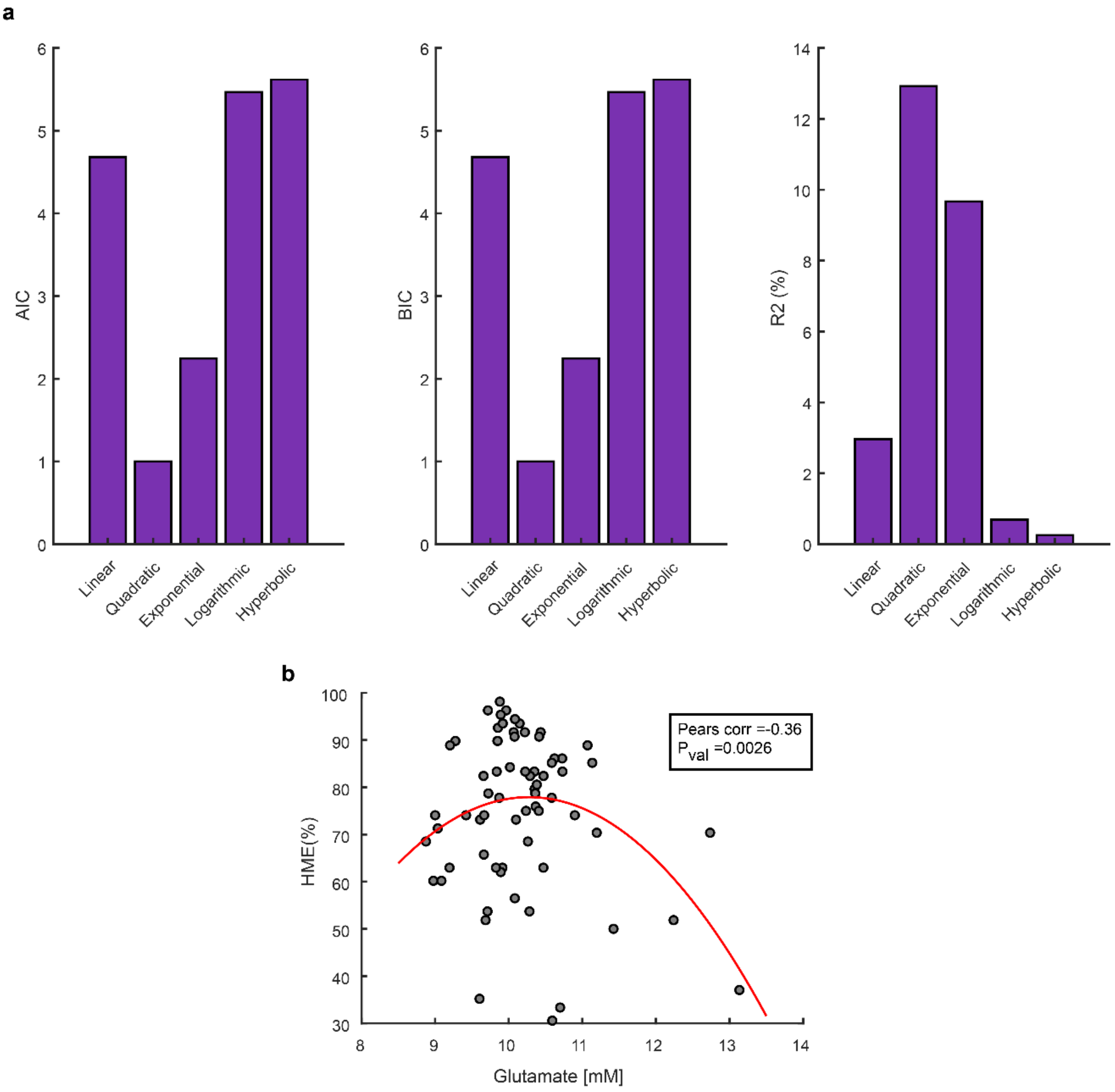


**Fig. S3: Investigating glutamate's relation with the percentage of high mental effort choices.** (**a**) Model comparison to explore the relationship between the number of high mental effort (HME) choices and glutamate concentrations. The quadratic model reaches both the lowest AIC and BIC score and results in the highest $R^2$ score, outperforming other models. (**b**) Pearson correlation between HME and centered and squared glutamate concentrations in the dmPFC/dACC (r =-0.36, p = 0.003).

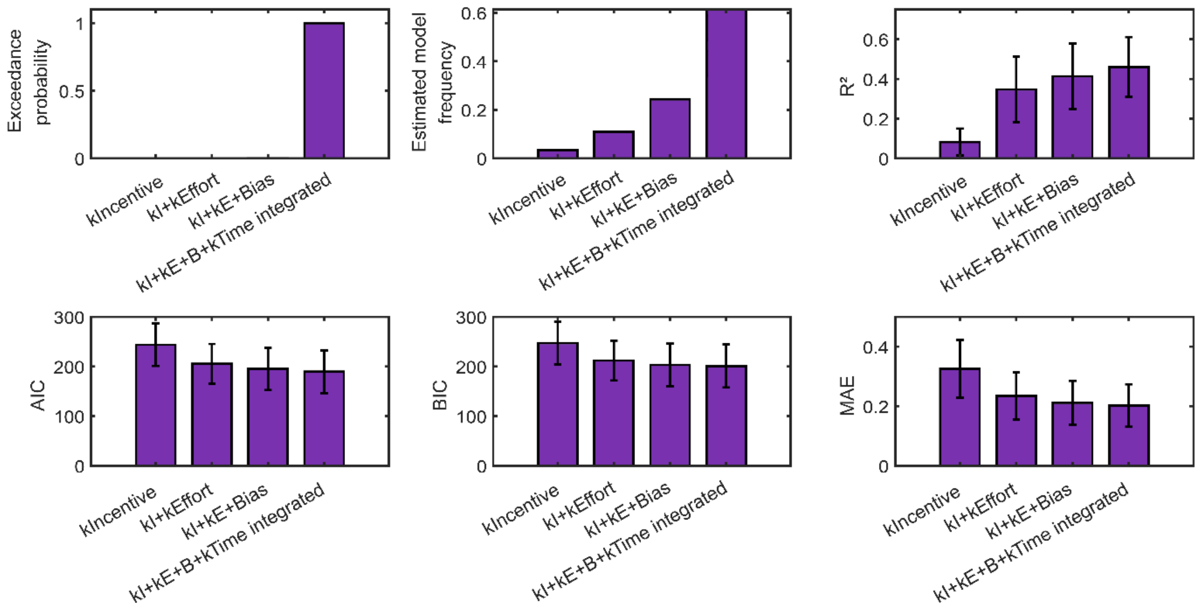


**Fig. S*4*: Model comparison between different behavioral computational models of effort-based decision-making in our task.** Confirmed unanimously by the exceedance probability score, estimated model frequency, Akaike information criterion (AIC), and Bayesian information criterion (BIC), our last model, including a sensitivity for effort, a bias term and integrating the effect of effort over time dynamically, resulted in the best fit. Furthermore, the winning model also resulted in the lowest mean absolute error (MAE), and the highest $R^2$ value, yielding the best description of the participant's decision-making.

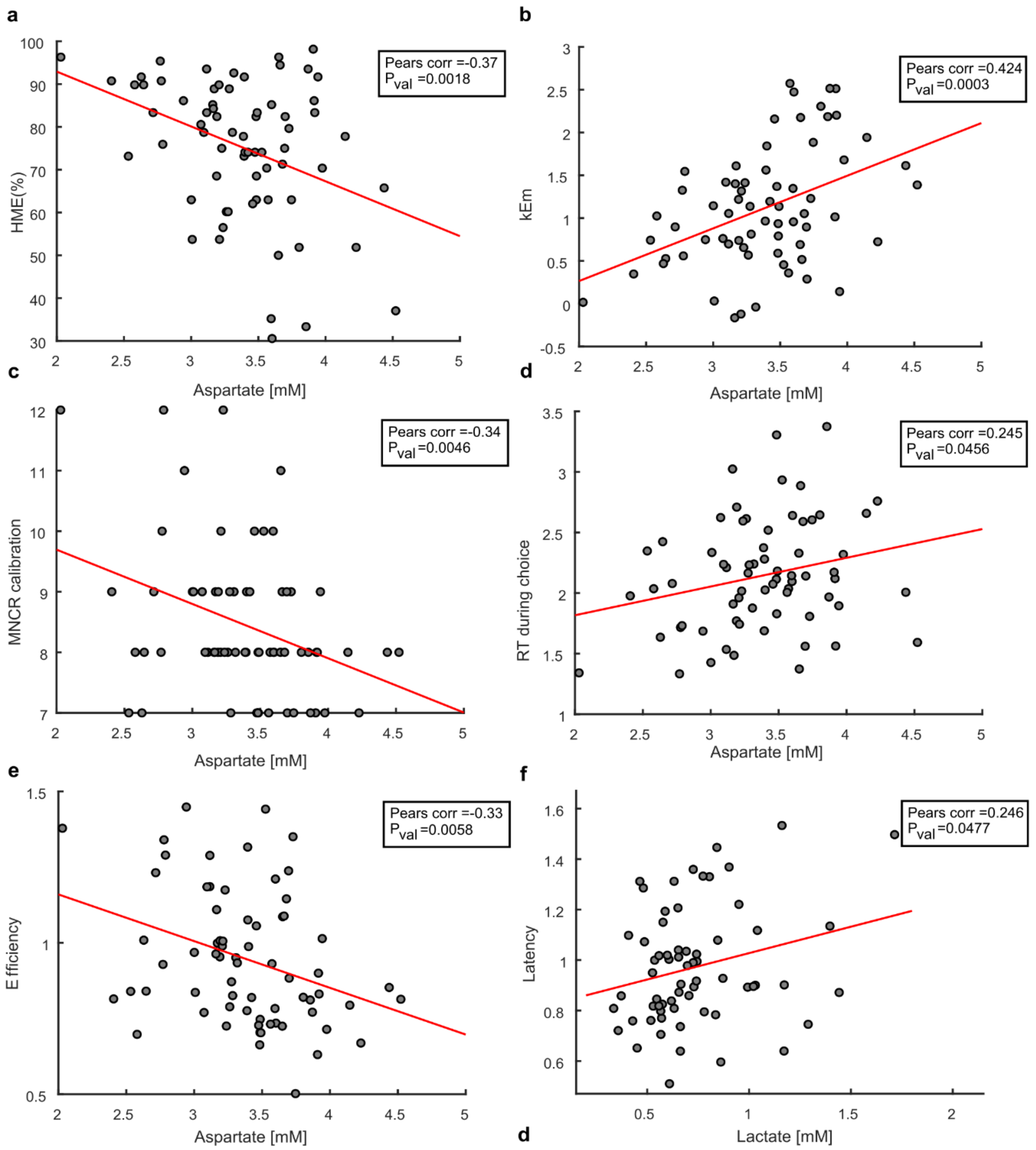


**Fig. S5: Correlation between measures of interest.** (**a**-**e**) Correlation between dmPFC/dACC aspartate concentrations and (**a**) HME choices ($r = -0.37$, $p = 0.002$), (**b**)mental effort sensitivity kEm ($r = 0.42$, $p < 0.001$), (**c**)minimum number of correct responses (MNCR) in the mental effort task (i.e. calibration scores) ($r = -0.34$, $p = 0.005$), (**d**) average choice reaction times (RT) across participants ($r = 0.25$, $p = 0.046$), and (**e**) mental effort efficiency ($r = -0.33$, $p = 0.006$). (**f**) Correlation between dmPFC/dACC lactate and the latency to start performing the mental efforts ($r = 0.25$, $p = 0.048$).

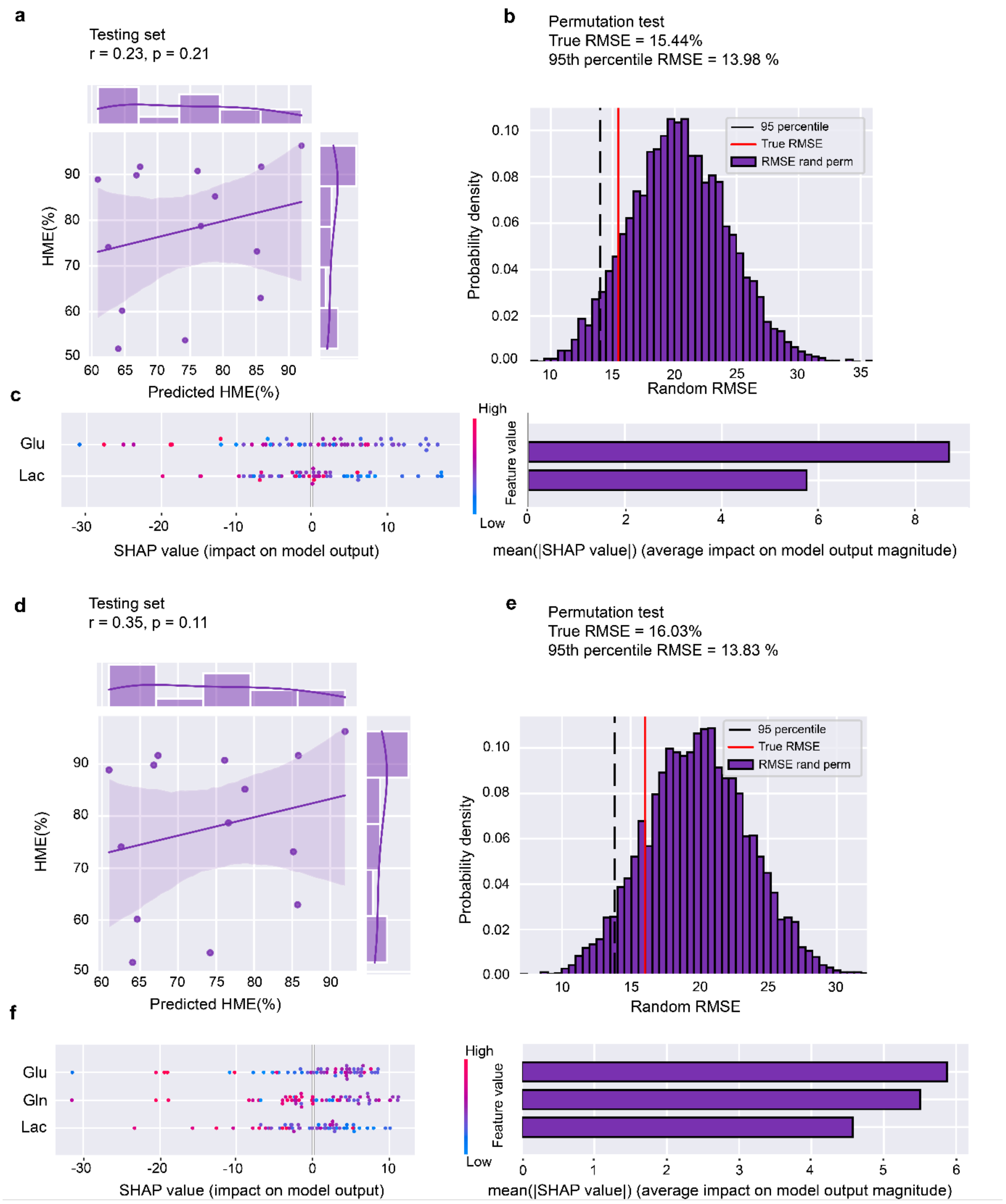


**Fig. S6: Model prediction of the percentage of high mental effort (HME) choices, based on metabolites concentration in the dmPFC/dACC, using only the combination of glutamate and lactate, or glutamate, glutamine, and lactate metabolites.** (**a**) Correlation between the XGBoost model's prediction and the percentage of times high mental effort (HME) was chosen for the test set (r= 0.34, p = 0.21). (**b**) Permutation test by permuting labels randomly and repeating the procedure, using the 95$^{th}$ percentile as a threshold (threshold 95$^{th}$ percentile = 13.98% < model RMSE = 15.44%). (**c**) SHAP values for the trained model were calculated for each subject. (**d**) Correlation between the XGBoost model's prediction and the percentage of times high mental effort (HME) was chosen for the test set (r= 0.35, p = 0.11). (**e**), Permutation test by permuting labels randomly and repeating the procedure, using the 95$^{th}$ percentile as a threshold (threshold 95$^{th}$ percentile = 13.83% < model RMSE = 16.03%). (**f**) SHAP values for the trained model for each subject.

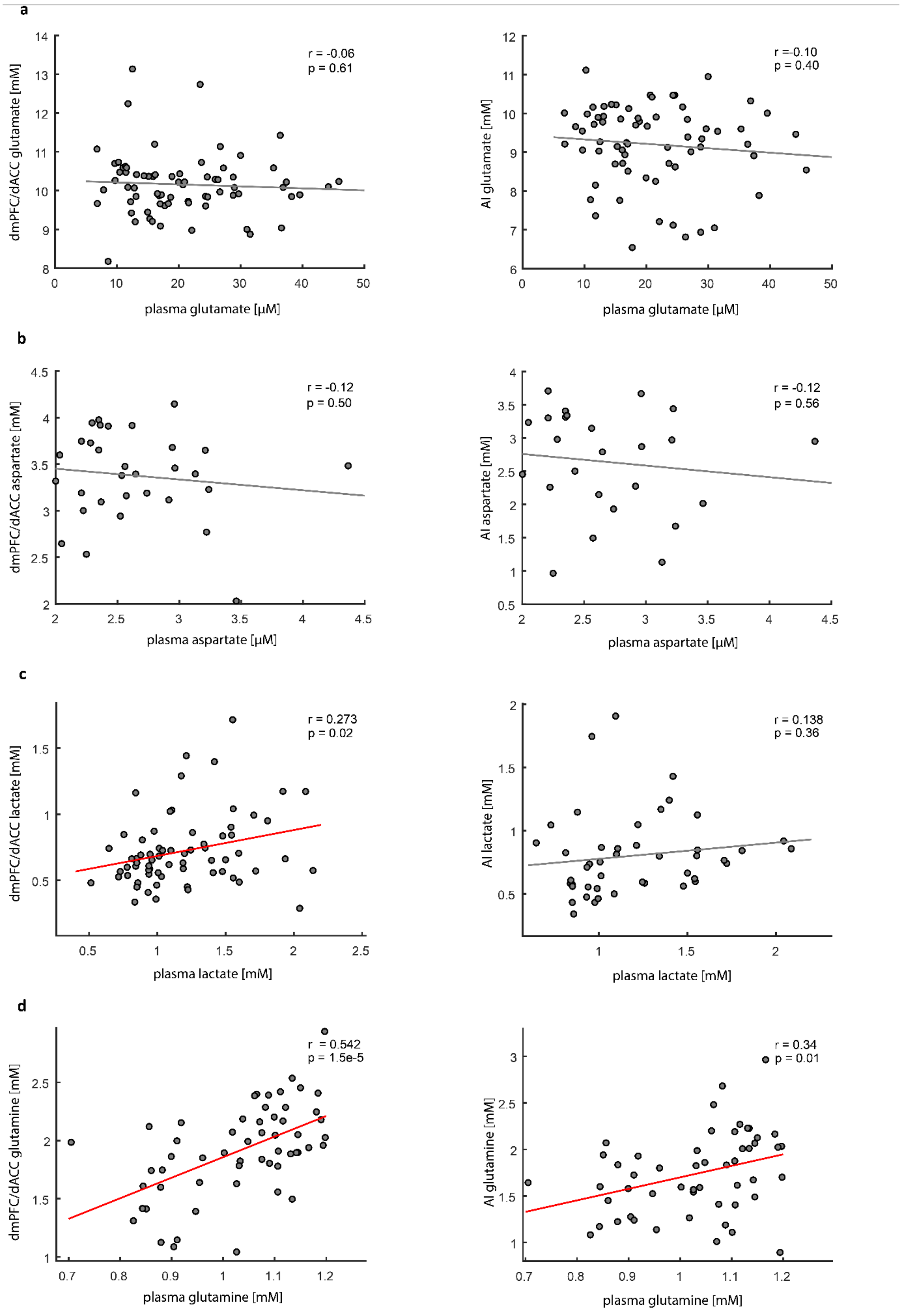


**Fig. S7: Cross-correlation of plasma and cerebral metabolite levels in the dmPFC/dACC and AI.** (**a**) Glutamate shows no significant correlation between plasma and brain concentrations in either region (dmPFC/dACC: r = -0.06, p = 0.61; AI: r = -0.10, p = 0.41). (**b**) Aspartate levels also display no significant correlation in both regions (dmPFC/dACC: r = 0.12, p = 0.51; AI: r = 0.12, p = 0.56). (**c**) Lactate shows a weak but significant correlation in the dmPFC/dACC (r = 0.27, p = 0.023) but not in the AI (r = 0.14, p = 0.36). (**d**) Glutamine exhibits a moderate to strong correlation, significant in both regions (dmPFC/dACC: r = 0.54, p < 0.001; AI: r = 0.34, p = 0.014), suggesting a more pronounced relationship between plasma and brain levels compared to the other metabolites.
